# OneNet – One network to rule them all: consensus network inference from microbiome data

**DOI:** 10.1101/2023.05.05.539529

**Authors:** Camille Champion, Raphaelle Momal, Emmanuelle Le Chatelier, Mathilde Sola, Mahendra Mariadassou, Magali Berland

## Abstract

Modeling microbial interactions as sparse and reproducible networks is a major challenge in microbial ecology. Direct interactions between the microbial species of a biome can help to understand the mechanisms through which microbial communities influence the system. Most state-of-the art methods reconstruct networks from abundance data using Gaussian Graphical Models, for which several statistically grounded and computationnally efficient inference approaches are available. However, the multiplicity of existing methods, when applied to the same dataset, generates very different networks. In this article, we present OneNet, a consensus network inference method that combines seven methods based on stability selection. This resampling procedure is used to tune a regularization parameter by computing how often edges are selected in the networks. We modified the stability selection framework to use edge selection frequencies directly and combine them in the inferred network to ensure that only reproducible edges are included in the consensus. We demonstrated on synthetic data that our method generally led to slightly sparser networks while achieving much higher precision than any single method. We further applied the method to gut microbiome data from liver-cirrothic patients and demonstrated that the resulting network exhibited a microbial guild that was meaningful in terms of human health.

## 1 Introduction

The human gut microbiota is a complex ecosystem consisting of trillions of microorganisms, mainly viruses, bacteria, archeae and microbial eucaryotes, that play critical roles in host physiology including digestion, immune function and metabolism Belkaid and Hand [2014], Chatelier et al. [2013]. Recent advances in sequencing technologies have enabled the characterization of gut microbiota composition and function at a fine scale, providing opportunities to understand the microbial communities that reside within the human gastrointestinal tract. However, despite these technological advancements, understanding the interactions within the bacteria of the gut microbiota remains a major challenge. These interactions are complex as microorganisms can interact with each other in a multitude of ways: through mutualism, parasitism, commensalism and competition to only cite a few Weiss et al. [2017], Faust and Raes [2016].

To address this challenge, network-based approaches have been developed to infer microbial interactions and construct microbial interaction networks. The resulting networks can reveal potential interactions between microbial taxa and support the identification of microbial guilds. Those guilds are defined as groups of microorganisms that co-occur and may interact with each other. Identifying microbial guilds is crucial for understanding the ecological dynamics of the gut microbiota and can provide insights into the role of the microbiota in health and disease Wu et al. [2021], Xiao et al. [2022].

Formally, microbial interaction networks consist of nodes, which correspond to microbial species, and edges, which correspond to interactions between those species. Positive and negative interactions are rarely observed directly. They are instead often reconstructed from abundance data, using either longitudinal data (see the generalized Lotka-Volterra model in Bucci et al. [2016]) or co-occurrence data. We focus here on the latter suite of methods.

The simplest way to identify microbial interactions is to perform a correlation analysis. However, correlation-based methods model total dependencies and are therefore prone to confusion by environmental factors (*e.g.* shared habitat preferences or susceptibility to the same abiotic factors) and do not lend themselves to a clear separation between indirect and direct effects Friedman and Alm [2012]. By contrast, conditional dependency-based method eliminate indirect correlations from direct interactions and lead to sparser and easier to interpret networks, at the cost of increased computational burden and more sophisticated models. The problem of network inference is complicated by the adverse characteristics of microbial abundance data, which are sparse, heterogeneous, heteroscedastic and show extreme variability. These data are thus tricky to model and leads to poorly reproducible and/or sparse networks, with many missing edges Peschel et al. [2021].

The most common framework for the estimation of the conditional dependencies is Gaussian Graphical Models (GGM) Lauritzen [1996], which describe the conditional dependency structure of multivariate Gaussian distributions. As microbiome abundance data don’t directly fit within the gaussian framework, three main workarounds are commonly used: data transformation, models based on alternative distributions and models based on latent variables; the whole strategy is also illustrated on Fig 1.

**Figure 1:**
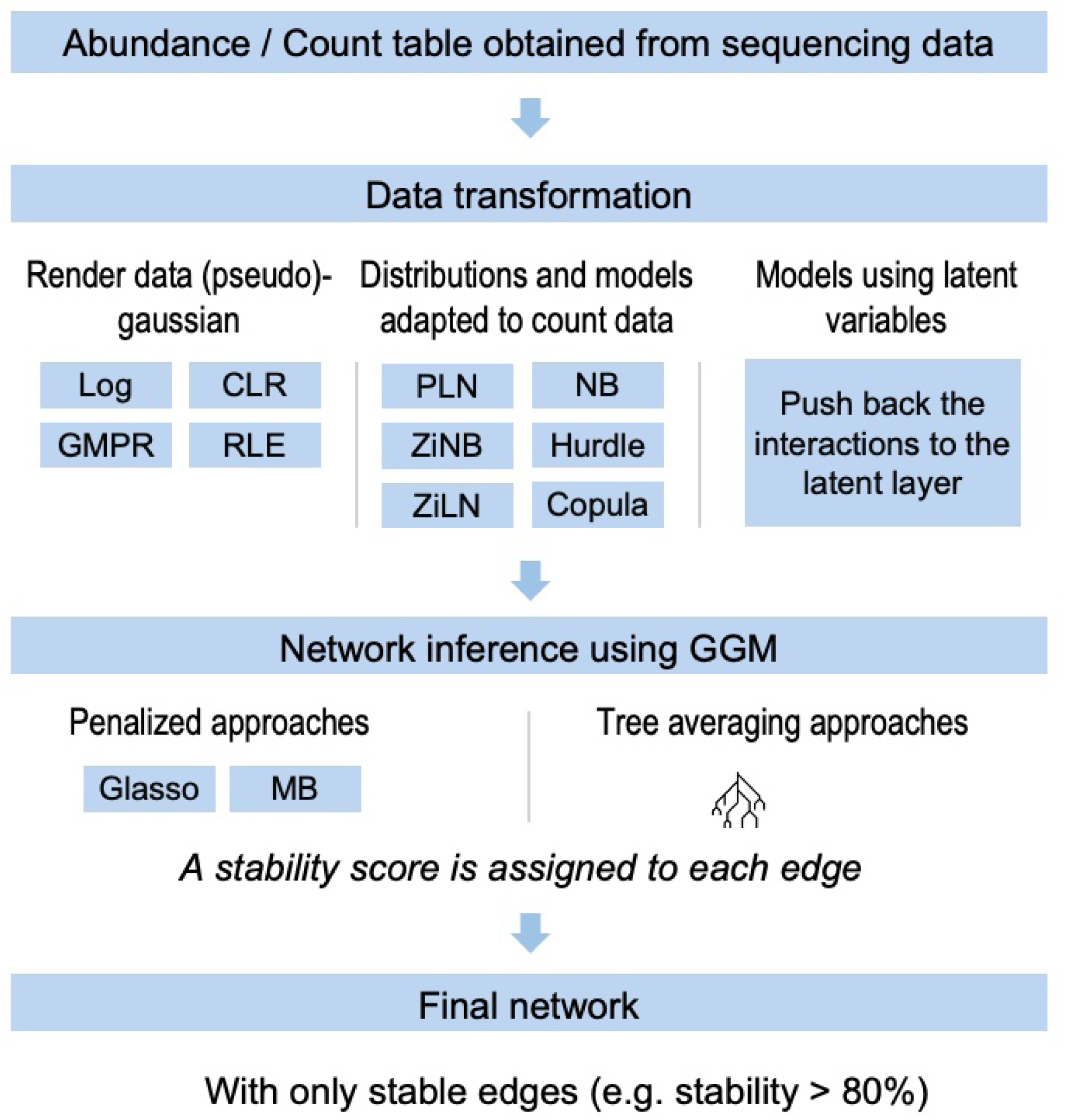
The classical network inference pipeline.

The complexity of reconstructing networks from co-occurrence data has spawned a rich literature with many methods relying on the solutions exposed above, including (i) approaches based on correlation as SparCC Friedman and Alm [2012], CoNet Faust and Raes [2016], (ii) approaches based on probabilistic graphical models as SpiecEasi Kurtz et al. [2015], gCoda Fang et al. [2017], SPRING Yoon et al. [2020], PLNetwork Chiquet et al. [2018], ZiLN Prost et al. [2021], COZINE Ha et al. [2020], Magma Cougoul et al. [2019], EMtree Momal et al. [2020] and (iii) approaches based on the inference of the latent correlation structure as CCLasso Fang et al. [2015], SparCC Friedman and Alm [2012]. The most recent methods include: mixPLN and ms-mixPLN Tavakoli and Yooseph [2019], Yooseph and Tavakoli [2022] which consider the problem of inferring multiple microbial networks (one per host condition) from a given sample-taxa abundance matrix when microbial associations are impacted by host factors. The HARMONIES approach Jiang et al. [2020] addresses some critical aspects of abundance data (compositionality due to fixed sampling depth, over-dispersion and zero-inflation of the abundances) while maintaining computational scalability and sparsity of the interaction network, in contrast to mixPLN and ms-mixPLN. Finally, NetCoMi Peschel et al. [2021], provides a one-step platform for inference and comparison of microbial networks, by implementing many existing methods for abundance data preprocessing, network inference and edge selection in single package.

All these methods have been designed to infer networks based on different mathematical hypotheses and thus have different strengths and weaknesses when modeling microbiome data. Each microbial network inference algorithm usually returns distinct edges to connect the taxa together, as many facets of the same reality. Methods to combine different microbial networks have been proposed before but they rely on spectral decomposition and do not work at the edge level Affeldt et al. [2016]. In this article, we present OneNet, an ensemble method that generate robust and reliable consensus network that will facilitate the identification of microbial guilds and generation of new hypotheses.

## 2 Methods

### 2.1 Overview of OneNet

We developed OneNet, a three-step procedure for robust consensus network reconstruction, illustrated on Fig 2.

**Figure 2:**
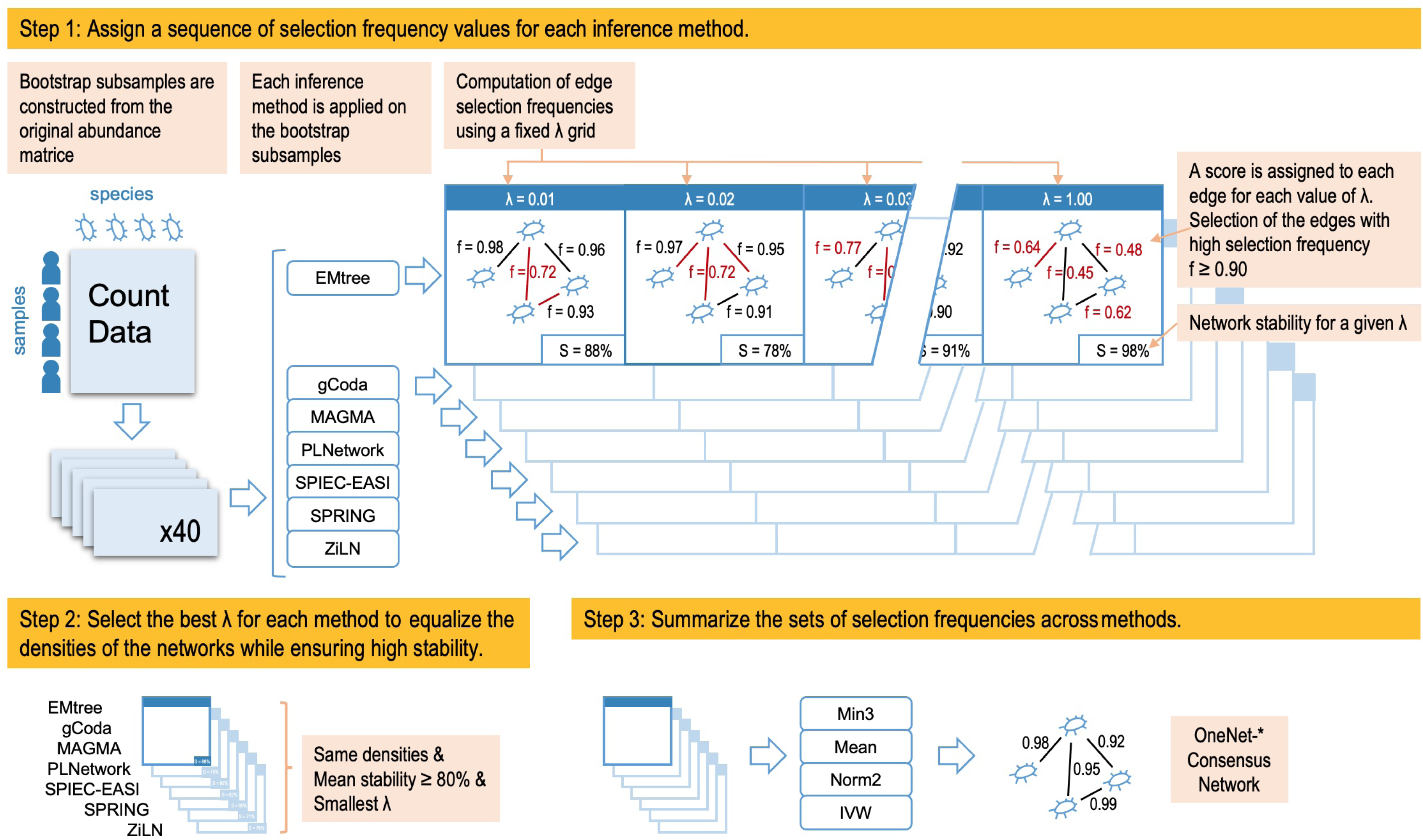
High level summary of the OneNet pipeline. (i) bootstrap subsamples are constructed from the original abundance matrice, (ii) each inference method is applied on the bootstrap subsamples to compute edge selection frequencies using a fixed *λ* grid, (iii) a different *λ* is selected for each method to achieve the same density in all methods, (iv) edge selection frequencies are summarized and (v) thresholded to compute the consensus graph.

We included seven inference methods in OneNet, all of which rely on Gaussian Graphical Models (GGM) to estimate conditional dependencies networks: Magma, SpiecEasi, gCoda, PLNetwork, EMtree, SPRING and ZiLN. We excluded mixPLN and ms-mixPLN as they do not reconstruct a single network but rather a collection of networks and NetCoMi as it collects already existing methods rather than introducing a new one. We left out HARMONIES from the comparison as its implementation doesn’t allow the user to specify the regularization grid, a crucial step in the ensemble method, and achieved worse performance than included methods in preliminary tests. We also excluded COZINE from OneNet because its implementation doesn’t rely on resampling and prevents it from being integrated, but we nonetheless included it in the benchmark as it compared favorably to others methods in preliminary tests. Table 1 summarizes the inference strategies adopted by each method and the potential integration of covariables in the model.

**Table 1:** Characteristics of the network inference methods.

| Method | Normalization | Distribution | Inference approach | Covariates | Reference |
| --- | --- | --- | --- | --- | --- |
| SpiecEasi | CLR | Multivariate gaussian | MB | No | Kurtz et al. [2015] |
| gCoda | CLR | Multivariate gaussian | glasso | No | Fang et al. [2017] |
| SPRING | CLR | Copulas | MB | No | Yoon et al. [2020] |
| Magma | GMPR + RLE | Copulas + ZINB | MB | Yes | Cougoul et al. [2019] |
| PLNetwork | GMPR + RLE | PLN + Latent variables | glasso | Yes | Chiquet et al. [2018] |
| EMtree | GMPR + RLE | Latent variables | Tree averaging | Yes | Momal et al. [2020] |
| ZiLN | CLR | Latent variables | MB | No | Prost et al. [2021] |
| COZINE | CLR | Hurdle gaussian | MB | Yes | Ha et al. [2020] |

Inference methods, abundances normalization (Centred Log Ratio (CLR), Geometric Mean of Pairwise Ratios (GMPR), Relative Log Expression (RLE)), distribution transformations, inference approaches (Meinshausen-Bühlmann (MB), glasso, tree averaging), covariates integration and references. OneNet is based on all methods but *COZINE*.

### 2.2 Step 1: Assign a sequence of selection frequency values to each edge for each inference method

Each inference method assigns a score to the edges: either a probability (for the tree averaging method) or the maximal penalty level *λ* above which the edge is selected in the network. An optimal penalty *λ^∗^* on these scores is then needed for an edge to be selected in the final network. Several approaches exist but the concept of stability selection Liu et al. [2010] is the most widely used and the one considered in this work as it yields a compromise between precision and recall, while fostering reproducibility. The associated method, called Stability Approach to Regularization Selection (StARS) uses a resampling strategy to select the value of *λ^∗^* leading to the most stable graph. We describe briefly in the following the StARS algorithm introduced by Liu et al. [2010] before presenting the modification of StARS we propose in this work.

#### 2.2.1 StARS algorithm (from Liu et al. [2010])

The full dataset *X* of size *n* is subsampled *B* times by selecting a subset of rows to create *B* subsamples *X*^(1)^*, …, X*^(^*^B^*^)^, each of size *n^′^* = 0.8*n* if *n ≤* 144 and 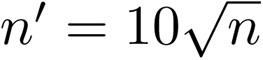 otherwise, following the recommendations of Liu et al. [2010]. The network inference is conducted on each subsample *X*^(^*^b^*^)^ for each value of *λ* in a grid (*λ*_1_*, …, λ_K_*) to obtain a graph *G^b,k^*, with *k* 1*, …, K* and *b* 1*, …, B*. The selection frequency of edge *e* for parameter *λ_k_*, is computed as its selection frequency across the subgraphs:

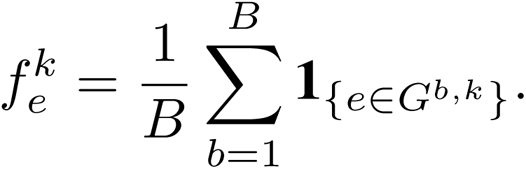

where 1_{a∈A}_ is the indicator function for *a* to belong to set *A*. The selection frequency over resamples gives an idea of edge reproducibility: frequency and robustness of the edges are clearly related. StARS aggregate those frequencies to construct a network-level measure of edge variability defined as:

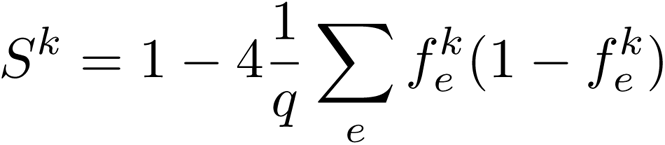

where *q* = *p*(*p* − 1)*/*2 is the total number of possible edges and *S^k^* can be thought of as the mean of (edge-level) Bernoulli variances. Each value *λ_k_* is associated to a single selection frequency vector, and a resulting stability value. Finding the right edge frequency is therefore equivalent to finding the right stability level. Classical choices for stability are *stab* = 80% or *stab* = 90% to have a good compromise between recall and precision and the optimal level *λ^∗^* is chosen as 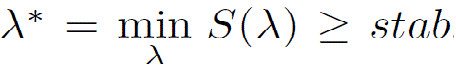. Once the optimal level *λ^∗^* is fixed, it is common practice to refit the model: forget the subsamples and run the inference method with the *λ^∗^* chosen by StARS on the full dataset *X* to return the corresponding graph.

#### 2.2.2 Using edge-level selection frequencies rather than network-level stability

We present here our suggested modification of the StARS algorithm allowing us to use edge-level frequencies to build consensus networks. Instead of computing the network-level stability *S^k^* and doing a final refit step, as is done in StARS, we instead select directly edges with high selection frequency to create the set 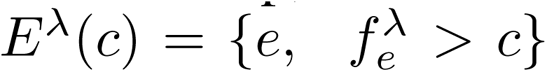 of highly reproducible edges, where *c* is a constant close to 1, typically 0.9 or higher. In this way, we guarantee both high precision and high reproducibility for edges in *E^λ^*(*c*) as they are selected many times in the resampling. Smaller values of *λ* give rise to larger sets *E^λ^*(*c*) and higher recalls. Two advantages of using frequencies rather than refitting the network are (i) filtering out edges with low support that could be included in the refit graph and (ii) making it easier to combine the edges inferred by the different methods.

### 2.3 Step 2: Select *λ* to equalize of the densities of the networks

In order to include the best edges in the consensus network, we must choose one *λ* per method. A natural choice would be the value 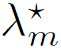 selected by StARS for the method *m*. However, we observed that StARS computes a very different precision/recall for each method as it does not integrate the same number of edges in the graph (Fig S12). We select instead the smallest *λ_m_* such that (i) the sets 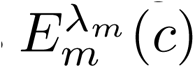 are roughly of equal sizes and (ii) the mean stability is above a given threshold: 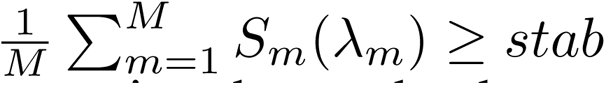. It forces all methods to contribute with a similar number of edges to the consensus while ensuring that each edge set is reproducible. In the following, we applied the mean-stability with a coefficient of 80% as mentioned in the previous subsection. In practice, to match edge set sizes, we worked with edge density rather than with *λ* values as the two are monotonically related.

### 2.4 Step 3: Summarize the sets of selection frequencies across methods

Building a consensus network from the sets of edges *E^λ^*(*c*) produced by the different methods is akin to an ensemble procedure where many methods are combined together.

In order to produce a stable and accurate consensus network, we define several summary metrics aimed at mitigating the drawbacks of each method while benefiting from their strengths. The consensus is obtained by summarizing edges frequencies across the methods. Denoting by *f_m_* the selection frequency of a given edge with method *m* 1*, …, M*, we define:

- *mean*: average selection frequency 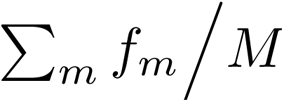,
- *norm2*: euclidean norm (2-norm) 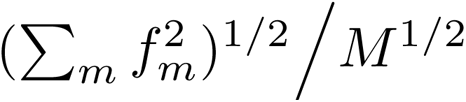,
- *IVW*: inverse-variance weighted average 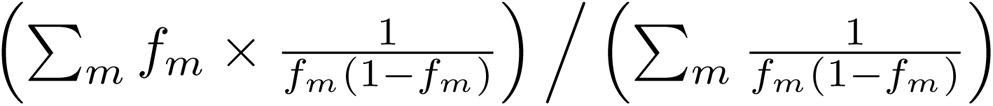, where *f_m_* follows a Bernoulli with 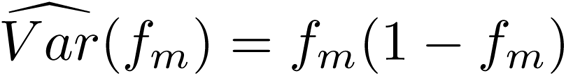,
- *minp*: high frequency for at least *p* methods 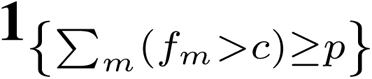.

We chose the mean as it is the most widely used metric to summarize a set of value, and *minp* as it is common practice in ensemble methods to keep a variable selected by at least *p* methods in the ensemble. Norm2 skews the summary towards high frequencies whereas Inverse-Variance Weighted (IVW) average upweights high and low frequencies as they are easier to estimate that mid-range frequencies. All summaries except *minp* can account for method-specific weights *w_m_* by replacing *f_m_* (resp. *M*) with *w_m_f_m_* (resp. 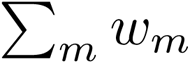). In this work, we only considered the simple case *w_m_* = 1. Finally, note that *minp* is the only one that returns a binary summary, all the other ones take value in [0, 1], like the original selection frequencies.

To reinforce the methodological assessment of OneNet, we also considered a naive consensus, called hereafter MRC for Majority-Rule Consensus, similar in spirit to min4 but applied directly to the refit networks, rather than to edge selection frequencies. MRC simply consists in running all methods with their default settings (*e.g.* without aiming for similar number of edges in all networks) and keeping only edges inferred in at least half the networks. Note that MRC and the different variants of OneNet have almost identical computational costs as all inference methods, but COZINE already include a StARS step.

### 2.5 Statistical analyses

In Figures Fig 4, S3 and Fig S4, all methods were compared in terms of PPV using a one-way ANOVA followed by Tukey’s HSD post-hoc test to assess pairwise significant differences. Results are shown using the compact letter display: two methods are significantly different if they don’t share a common letter and small letters (*e.g.* “a”) correspond to methods with higher PPV. Boxplots use the following convention: the box ranges from the first (Q1) to the third (Q3) quartile with the median (Q2) correspond to the line within the box, whiskers extends 1.5 × (Q3 – Q1) below and above the box and all points outside the whiskers are considered outliers and shown as dots.

### 2.6 Datasets

#### 2.6.1 Simulated dataset

In this work we simulate data using the methodology described in Yoon et al. [2020] which is based on gaussian copula to control the network structure followed by sampling from the species marginal distributions to preserve the peculiarities of abundance data. This method yields synthetic data with marginal distributions that are closer to the original empirical dataset, while enforcing a given correlation structure between the species.

To generate the simulated dataset, we use in input the empirical dataset described below restricted to diseased individuals and transform the abundance table into count table.

The dataset is simulated in the framework of an unknown undirected graph *G*(*V, E*), with no retroactive loop, consisting of *p* vertices *V* = *{*1*, …, p}* and a set of edges *E ⊆ V ×V* connecting each pair of vertices. The graph G is represented by its adjacency matrix *A* = (*A_ij_*)_(_*_i,j_*_)_*_∈E_* of size *p × p*, defined as:

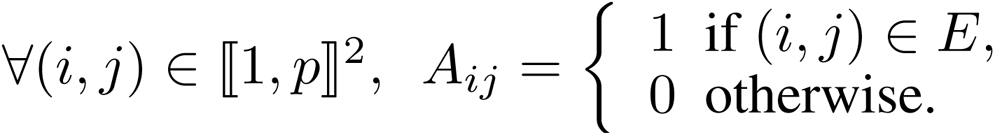

The package EMtree v.1.1.0 Momal [2021] is used to generate a precision matrix Ω defined as the graphical Laplacian *A* of a cluster graph. Ω is inverted to create the correlation matrix Σ and the idea was then to simulate variables with arbitrary marginal distributions from multivariate normal variables with correlation structure given by Σ using gaussian copula. Specifically, we generate a *n × p* matrix *Z* with independent normal rows 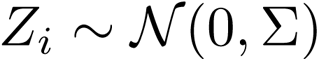. We then get uniform random vectors by applying standard normal cdf transformation to each column of 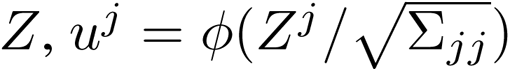 element-wise, and finally apply the quantile functions of the empirical data marginal distributions to each *u^j^*. The function synthData_from_ecdf from the SPRING package v.1.0.4 Yoon [2022] is used for these simulation steps. To assess the effect of sample size, we simulate datasets of size *n ∈* {50, 100, 500, 1000}.

### 2.7 Evaluation criteria

Each method is evaluated by comparing the inferred network structure to the known simulated network structure using the following metrics:

- Precision (positive predictive value): PPV= TP/(TP+FP),
- Recall (true positive rate): TPR= TP/(TP+FN),

where TP stands for True Positive (a correctly detected edge), FP for False Positive (an edge detected where none should be) and FN for False negative (an undetected edge). The precision measures the proportion of real edges among the detected ones, whereas the recall measures the proportion of real edges which are detected.

#### 2.7.1 Empirical dataset

The empirical dataset, studied in Qin et al. [2014], corresponds to stool samples from 216 Chinese individuals sequenced using whole-metagenome sequencing techniques. The raw sequences are available as BioProject PRJEB6337 in the European Nucleotide Archive (ENA). Among this population, 102 individuals are healthy and 114 suffer from liver cirrhosis. Abundances of all microbial species (metagenomic species or MSP) detected using 10.4 million IGC2 gut gene catalogue Wen et al. [2017] are extracted using the Meteor software suite that creates a gene abundance table by mapping high quality reads onto the gene catalogue, using Bowtie2. Abundance of each MSP is computed as the mean abundance of 100 marker genes selected for that MSP, where the gene abundance is the read abundance normalized by the gene length. The abundance table is then transformed into a count table of size 1990 MSP by 216 individuals Champion et al. [2023]. In the Application section, we used only the 114 cirrhotic patients.

## 3 Results

In this section we evaluated the performances of both OneNet and the network inference methods on the simulated dataset.

### 3.1 Influence of the stability level on the inferred graphs

We first evaluated the effect of stability level on the performance of the inference methods. Instead of fixing a target stability at 0.8 or 0.9, we studied the relationship between the precision and recall of the inferred edges by each method for different stability levels. Because interactions between highly prevalent species are easier to reconstruct, we only kept metagenomic species with a prevalence greater than 50% (159 species). We let the sample size vary in {50, 100, 500, 1000} and we considered *B* = 40 resamples each time.

#### Methods have distinct precisions for a given stability level

Fig S1 shows the relationship between the performance obtained with the edge set *E^λ∗^* (0.90) (precision PPV90 and recall TPR90), and the corresponding stability. The difference in patterns grows with sample size *n*, revealing peculiarities inherent to each method. Clearly, methods have distinct performances for the same stability level. We observed that methods cluster in groups (glasso, neighorhood selection, tree aggregation) with different precision/recall tradeoffs. As a result, they produce edge sets that greatly differ both in size and quality. This suggests that the stability value is not a good indicator of the precision level achieved by each method.

#### Methods have comparable precision and recall for a given density level

Unlike precision, which is unavailable when dealing with empirical datasets, the density, or number of detected edges, can always be computed. Fig S2 shows the relationship between precision (resp. recall) and density for all methods at increasing sample sizes. The curves are almost superimposed for values of *n* up to 100, after which different behaviors appeared. However, the gap in performance between methods stayed small when imposing the same density, rather than the same stability. This also meant that, whatever the method used, the *m* first edges included in the graph achieve similar graph reconstruction quality, although they correspond to different stabilities. We thus selected individual graphs based on density, rather than stability, to include only graphs with similar precision and recall in the consensus phase.

#### Mean stability as a proxy of the density level

The previous observation prompted us to explore the link between density and stability for different values of *λ*. Fig S3 shows how stability decreases with increasing density. We set a target mean stability value (e.g. 90%) for each value of *n* (here between 50 and 1000). As *n* grows, we observed that the density increased from 90 to 170, as well as the spread between methods. For *n* = 1000, stabilities ranged from 0.8 for EMtree to 1 for SPRING. We can see how targeting the mean stability rather than the same stability for all methods allows to adapt the precision level of each method through density to make them more similar.

#### Mean stability increases the consistency between the performances of the network inference methods

We compared, for different sample sizes, the precision – recall tradeoff achieved by the mean stability to the ones achieved by a fixed stability (e.g. 0.9). Figure 3 shows that the sample size has a major impact on the precision and stability. The ROC curves stabilize to near-perfection starting from *n* = 500 (Fig 3c). It is also noticeable that the adapted stabilities reduced the range of the method’s precision. Furthermore, for glasso-based methods (gcoda, PLNnetwork), the new target led to a 20 points improvement in TPR at almost no cost in PPV, for large sample sizes. Note that COZINE performs especially poorly compared to other methods for large sample sizes (*n* = 1000, Fig 3d), with a TPR close to 1 but a PPV close to 0. This is due exclusively to the use of a fast BIC critera instead of the resampling-based StARS which selects a small *λ* leading to a very dense graph in this example. Very poor performances of COZINE are also observed on large sample sizes (*n ≥* 500, Fig S10 and Fig S11) but not on small ones (*n* = 100, Fig S9).

**Figure 3:**
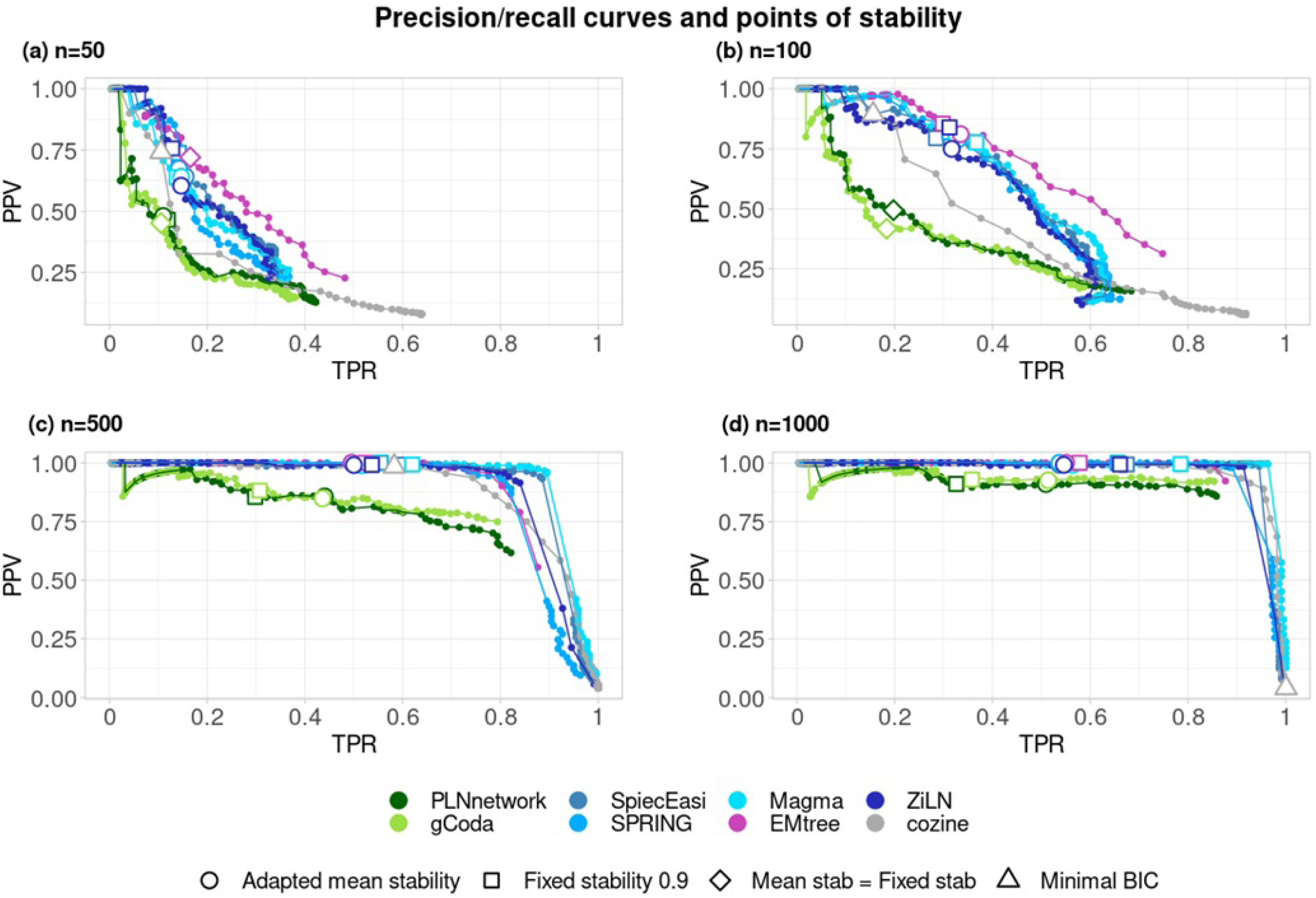
Precision – recall curves of each inference method for different sample sizes. (a) 50 (b) 100 (c) 500 (d) 1000. The TPR/PPV compromise achieved for *λ^∗^* corresponding to a stability of 0.9 is shown with a circle, the one achieved by a mean stability across methods of 0.9 is shown with a square. Whenever the selected *λ* is the same, the circle and the square are replaced with a diamond. Finally, note that COZINE relies on minimization of a BIC criteria (shown with a triangle) rather than on the resampling-based stability selection to choose the regularization parameter.

### 3.2 OneNet versus the classical network inference methods

We computed, for a frequency threshold of 90%, the precision and recall values obtained by the classical network inference methods (COZINE, gCoda, PLNnetwork, SPRING, Magma, SpiecEasi, ZiLM and EMtree) and we compared them to the OneNet networks (with the summay metrics mean, norm2, IVW and min3). Note that because of the similar density, each method provided roughly the same number of edges to OneNet. Fig 4 shows that OneNet with the mean and norm2 consensus methods, achieved the best precision levels (one-way ANOVA followed by Tukey’s HSD post-hoc test, *p <* 2.10*^−^*^16^) but the worse recall values. OneNet with the min3 summary has comparatively lower precision but higher recall and OneNet with the IVW summary has both worse recall and worse precision. This illustrates how OneNet generally leads to sparser networks with higher precision.

We also observed that the MB-based ones tend to outperform glasso-based ones. However, OneNet still demonstrated precision levels equivalent to the best methods when including (Fig S4) or not (Fig S5) the methods based on graphical lasso. This reflects the inherent robustness of consensus measures to methods with outlying performance.

**Figure 4:**
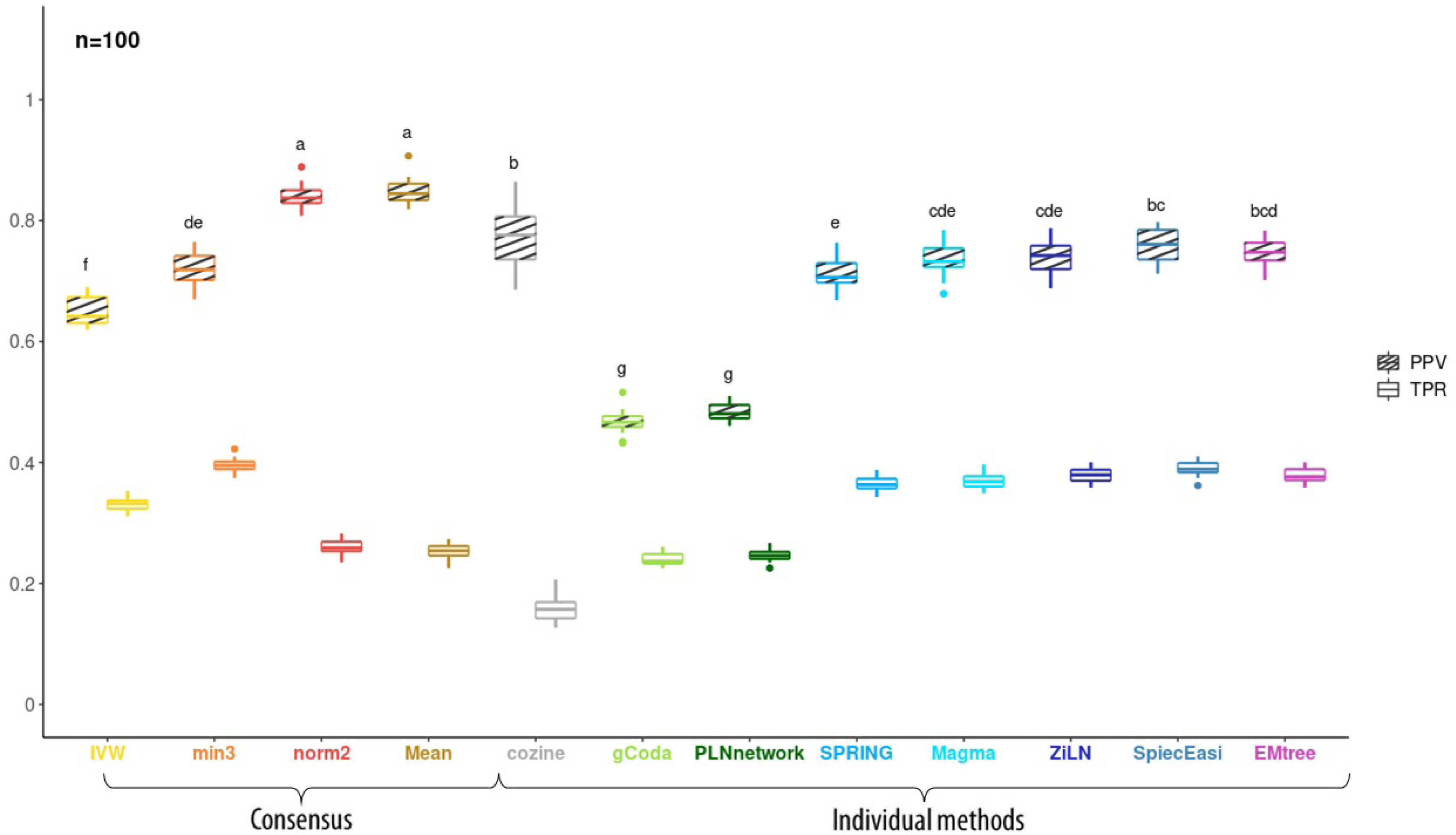
Quality of both single-method and consensus network in terms of PPV/TPR. assessed on simulated datasets of size *n* = 100 samples. Striped (resp. no-strip) boxplots show PPV (resp. TPR) values.

On top of that, sample size *n* has a dramatic effect on all criteria and methods. For large sample sizes (*n* ≥ 500), most methods exhibit precision above 99% and recall between 30% and 60% (Fig S4 and Fig S5, panels (c) and (d)). By contrast (Fig S4a and Fig S5b) for smaller but more realistic sample sizes like *n* = 50 (resp. *n* = 100), the median precision drops below 60% (resp. 80%) for all methods except OneNet-Mean and OneNet-norm2 which both remain above 75% (resp. 85%). Likewise, the recall drops below 40% for *n* = 100 (Fig S5b) and below 20% for *n* = 50 (Fig S5a).

We observed a discrepancy in terms of precision and recall for the COZINE method between Figure 3 and Fig S5. We hypothesized that it’s due to the original COZINE procedure (BIC criteria) used to select the optimal network, which leads to a dense graph (very high recall, very low precision, and therefore many spurious edges, see Figure 3). By contrast, Fig S5 shows the precision and recall values obtained with the resampling approach applied to COZINE are in line with other methods. This is an extreme example of lack of robustness, where the network reconstructed from the full dataset differs drastically from the ones reconstructed on random subsets of the data and illustrates the benefits of combining the resamples rather than doing a refit. Note that the difference between refit and combined resamples is much smaller for other methods as they already include a stability selection step designed to mitigate those problems. To explore further the benefits of combining edge selection frequencies rather than edges in the refit models, we compared OnetNet variants to the MRC consensus (Fig S13). We observed that MRC achieves a slightly higher TPR but a slightly lower PPV than OneNet-mean and OneNet-norm2 on synthetic data and thus constitutes a relevant approach.

Finally, the upset plots of edges detected by the inference methods for *n* = 100 (Fig S7) and *n* = 500 (Fig S8) show the same trends: (i) a large, but decreasing with *n*, number of edges undetected by any method, (ii) most reconstructed edges (whether wrongfully or not) detected by many methods and (iii) very few edges detected by only one or two methods (with the exception of gCoda and PLNnetwork). This strong agreement between all methods and their overall very high precision on large datasets (*n >* 500) is likely a consequence of our semi-parametric simulation setting and may constitute an optimistic evaluation of the methods.

## 4 Application to liver cirrhosis

To investigate the performance of OneNet relative to the other methods, we inferred all the networks from the microbiome dataset of cirrhotic patients presented in the Methods Qin et al. [2014]. Because of the small size of the dataset (114 samples), only metagenomic species with a prevalence greater than 50% were kept (155 species). The networks have been clustered using the CORE-clustering algorithm to reconstruct microbial guilds Champion et al. [2021]. Following the guidelines of this paper, we fixed the number of clusters between 10 and 19.

Fig 5 illustrates the inferred and clustered networks. We note that the OneNet-mean network is the sparsest one, consistent with our results on the simulated datasets (lower TPR but higher PPV as shown in Fig. 4 and sparser graphs as shown in the upset plots Fig S7 and Fig S8). Applying CORE-clustering to each network always resulted in 3 to 4 guilds.

**Figure 5:**
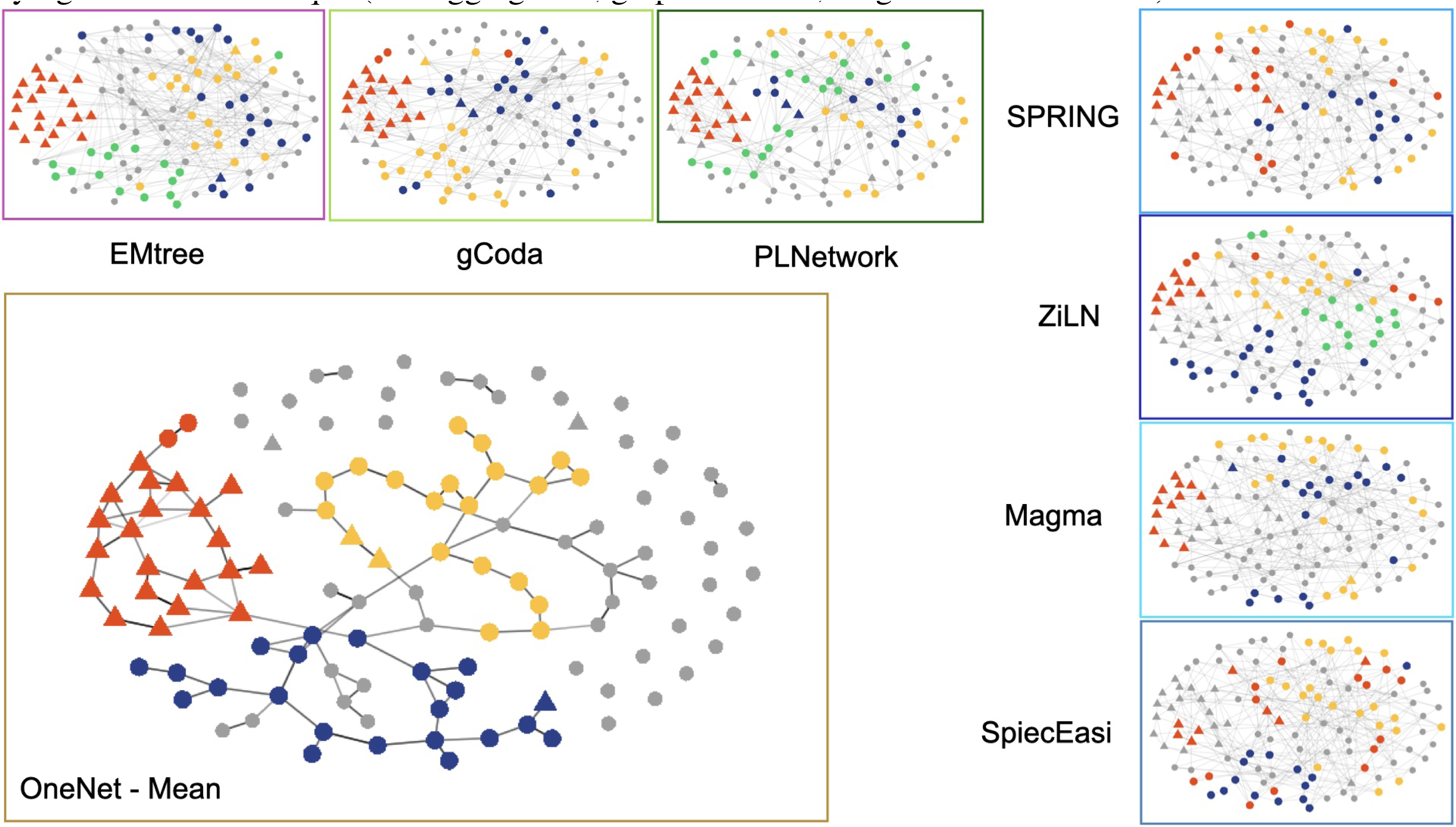
Consensus and single-method networks inferred on the liver cirrhosis dataset, followed by CORE-clustering algorithm to identify the microbial guilds. All graphs share the same layout, computed on the OneNet-mean network, to ease comparisons. Nodes are colored by cluster, with red always used for the cirrhotic guild in all graphs where it’s (at least partially) recovered and oral species are represented by a triangle. Methods are grouped based on the underlying inference technique (tree aggregation, graphical lasso, neighborhood selection).

One guild – the “cirrhotic guild”, shown in red in Fig. 5 and highlighted in Fig 6, is notable as it contains species associated to chronic diseases: obesity after weight-loss intervention (OB after WL) Liu et al. [2017], schizophrenia (SCHIZO) Zhu et al. [2020]), atherosclerotic cardiovascular disease (AVCD) Jie et al. [2017], Crohn disease (CROHN) He et al. [2017] and liver cirrhosis (LIVER) Qin et al. [2014] (Tab S1). Interestingly, the majority of the species detected in the cirrhotic guild, represented by a triangle shape, are oral. This result fits with Qin et al. [2014], which highlights the invasion of oral species in liver cirrhosis.

**Figure 6:**
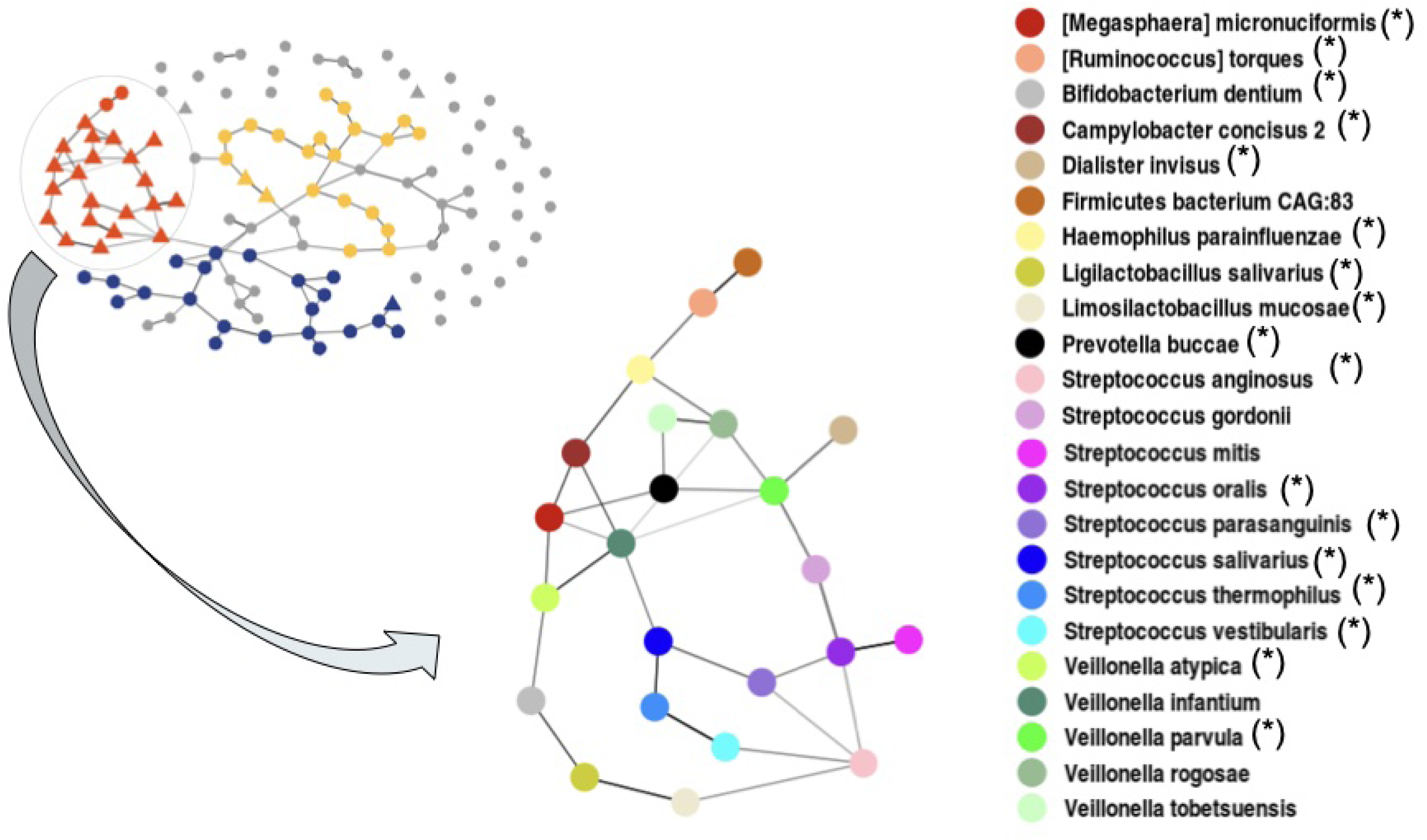
Detailed view of the cirrhotic guild identified in the OneNet-mean network with taxonomic information on the nodes. All species known to be associated with chronic diseases are marked with a star (*).

Among the 23 identified species of the cirrhotic guild, 17 have been found associated together in networks of chronic diseases (see Tab S1). These results are therefore consistent with what has already been shown in the litterature.

Among the different methods included in the consensus, only some (MAGMA, gCoda, PLNnetwork, EMtree, ZILN) were able to recover (at least partially) the cirrhotic guild from the consensus (highlighted in red in Fig. 5). The upset plot Fig S6 also shows greater disparities between methods and greater differences between norm2– and mean-consensus (with mean being the more restrictive) than expected from the synthetic data. In particular, some neighbordhood selection methods (SPRING, SPIEC-EASI, MAGMA) and EMtree have a lot of unique edges (Fig S6), in contrast with their behavior on synthetic data (Fig S7 and Fig S8) suggesting that they react differently to model misspecifications. Note that all methods selected by the mean-consensus are selected in at least 5 (out of 7) individual methods showing that mean-consensus is more stringent than a simple majority rule.

By contrast, the network and microbial guilds reconstructed from the healthy individuals (*n* = 102 individuals and *p* = 151 taxa with prevalence higher than 50%) using the same parameters exhibited a completely different structure (Fig S14). While we recovered the same number (3) of guilds, none matched the cirrhotic microbial guild. Most taxa from the the cirrhotic guild were either not present in the “healthy individuals”-network (as they didn’t pass the prevalence threshold in the healthy individuals dataset) or had much fewer interactions and were not included in a guild. Overall, this suggests that the interactions identified between the cirrhotic species are context-dependent, a finding that has already been documented in other contexts Weiss et al. [2023] and K. Faust [2021].

## 5 Discussion

The proposed framework, with a microbial consensus network inference method, offers new insights about inferring robust and sparse microbial networks. OneNet is robust in the sense that i) it uses GGM adapted to deal with the peculiarities of microbial abundance properties (inclusion of environmental effects as covariates, stabilization of data variability, adaptation to abundances with high proportion of zeros, etc), ii) it depends on seven network inference methods aiming for sparse and reproducible microbial network using either glasso, neighborhood selection or tree averaging approaches, iii) it relies on a three-steps procedure to improve the precision and reproducibility of both inference methods and OneNet. Indeed, the selection of edges with high inclusion frequencies and harmonization of stability selection achieve similar precision levels accross methods. The resulting consensus network uses a summary of the edge inclusion frequencies.

Results from the studies on synthetic and real data illustrated the first major and reassuring fact, that the inference methods overall agree with one another and with a fraction of the truth (Fig S7 and Fig S8), the remainder being hard-to-reconstruct edges. It then showed the effectiveness of OneNet compared to the inference methods. Among the different summaries considered, the mean or norm2 are preferred since they lead to slightly sparser networks but achieved much higher precision than any inference method, especially for sample sizes around *n* = 100, which is typical in microbiome studies. By contrast, min3 and IVW summaries gave a significant additional quantity of edges compared to the other summary metrics, yielding TPR levels that are comparable to those obtained with glasso-based methods without increasing the PPV, especially when the number of samples is small (*n ≤* 100) (Figures Fig S6).

In all numerical experiments, we showed that a minimal sample size to maintain high robustness was *n* = 100. In this scenario we suggest to use the mean summary in OneNet, as it proved to be more robust to small sample sizes. Obviously, the precision is affected by both the number of samples and microbial species in the system, the latter being controlled by the prevalence threshold imposed at the very beginning of the analysis. As illustrated on Figures Fig S9, Fig S10 and Fig S11, the prevalence threshold can be adjusted to increase the precision of the method depending on the number of samples (*prev* = 0.50 for *n* = 100 and *n* = 500, *prev* = 0.20 for *n* = 1000). From *n* = 1000, when considered individually, the neighborhood selection and the tree averaging approaches showed performances that were similar to OneNet. In this context, it could be possible to select one of these three approaches.

We also investigated a simple alternative (MRC) based on majority rule for edge selection. This simple alternative was competitive to OneNet variants on synthetic data, with slightly lower PPV and slightly higher TPR. We however observed that the inference methods produces results that were more contrasted in terms of edge sets on real data than on synthetic data (compare Fig S6 and Fig S7). It means that inference methods react differently to model misspecification in real life settings and may have lower TPR and PPV than estimated from the simulations and that some methods may contribute with a large number of spurrious edges to the consensus. This is for example the case of COZINE for large sample sizes in our simulations. While this shortcoming also applies in principle to OneNet variants, it is mitigated by using roughly the same number of edges from each method (step 2 of the algorithm). Finally, MRC is sensitive to methods which can fail drastically, as COZINE, whereas the compulsory resampling step used in OneNet stabilizes the results of each methods and prevents catastrophic behavior. This is less of an issue if all methods of the consensus use stability selection but then the computational overhead of MRC and OneNet variants are comparable. Overall, both OneNet and MRC perform better than no consensus but, compared to MRC, OneNet(-mean) is more robust than MRC and favors PPV to TPR.

An advantage of OneNet is its ability to easily incorporate new inference methods as soon as they are amenable to the modified stability selection framework used here. This is the case for all the methods considered in this work but COZINE. Indeed, COZINE relies on the BIC criteria to tune the regularization parameter *λ*: it doesn’t allow the user to provide a fixed *λ* grid for comparisons with other methods and doesn’t produce the table of edge selection frequencies required to compute summaries. This is however due to implementation choices (using BIC instead of StARS for selecting *λ*) rather than to fundamental incompatibilities with the OneNet framework.

## 6 Supplementary Methods

We detail here each step of the whole network inference strategy, illustrated on Fig 1. As microbiome abundance data don’t directly fit within the gaussian framework, three main workarounds are commonly used: data transformation, models based on alternative distributions and models based on latent variables:

### Transformations

A small constant is added to each abundance before log-transforming them. However, this transformation does not stabilize data variability because the log-transformed abundances scale with sequencing depths and covary with it, making dependencies modeling tricky. On the contrary, the Centered Log Ratio (CLR) transformation [Aitchison, 1982] guarantees the study of dependencies. It is however highly criticized when data contain a high proportion of zeros: proportions higher than 90% are typical in whole-metagenome or amplicon sequencing data. To circumvent this problem, Yoon et al. [2020] introduced a modified version of the CLR transformation (mCLR) that respects the original ordering of the data but doesn’t account anymore for the compositional nature of the data, the primary motivation of the CLR transformation. An alternative to compositional transformations is to use a normalization factor, such as Geometric Mean of Pairwise Ratios (GMPR) [Chen et al., 2018], Relative Log Expression (RLE) [Anders et al., 2010] and others like Wrench normalization factors [Kumar et al., 2018] or Cumulative Sum Scaling [Paulson et al., 2013]. The GMPR normalization is designed for abundances with a high proportion of zero values. It compares pairs of samples based only on the species they share, and considers the geometric mean of the median ratio. This makes this technique robust to both differentially abundant species and extreme values. When two samples do not share any species, the computation of GMPR fails (this happens when samples come from very contrasted conditions with no or limited species overlap). In this case, Relative Log Expression (RLE) normalization method can be used. This method is based on the assumption that most of the species are not differentially abundant. However, this normalization factor fails when no single species is shared across all samples, which is frequently the case in microbiome data. A modified version of RLE only considers positive abundances to avoid this drawback.

### Distributions and models

The second workaround is to use models adapted to abundance data characteristics: overdispersion (excess of variability in the data) and zero-inflation (excess of zeros). The **Poisson-log normal (PLN) model** [Chiquet et al., 2021] is designed for the analysis of abundance tables. It accounts for both structuring factors and potential interactions between the species. In the presence of overdispersion, the Poisson regression model is not adequate and can lead to biased parameter estimates and unreliable standard errors estimates. The **Negative Binomial (NB) model** is then often used [Forbes et al., 2010]. Both models can be seen as compound Poisson model (with a lognormal for the PLN distribution and Gamma for the NB) that are overdispersed compared to a *vanilla* Poisson distribution but the PLN is multivariate and can account for correlations between abundances. Contrary to the NB model, the **zero-inflated model** [Greene, 1994] is often motivated by an excess of zeros in the data, but less flexible than the zero outcome model. An intuitive approach to analyzing zero-inflated abundance data is to view the data as arising from a mixture distribution of a point mass distribution at zero and an abundance distribution. **Hurdle models** [Cragg, 1971] are a class of models for abundance data that help handle excess zeros and overdispersion. In contrast to Zero inflated-models, hurdle models capture both an excess or a lack of zeros in the dataset. The **zero-inflated negative binomial (ZINB) model** [Cheung, 2002], obtained by applying ZI to NB model, takes into account both overdispersion and excess of zeros. Finaly, **copulas** are a multivariate cumulative distribution functions for which the marginal probability distribution of each variable is uniform on the interval [0, 1]. As they fully describe the dependency structure, models with copulas allow to separate the modeling of marginal distributions (*e.g.* overdispersed, with excess zeros, etc) from the modeling of dependencies. Recent developments used gaussian copula coupled with arbitrary discrete marginal distributions to study multivariate abundance data [Anderson et al., 2019]. Popovic et al. [2019] showed that Gaussian copulas are a relevant and promising approach to the problem of network inference from abundance data, even if the computational cost is higher than for other methods. One way of taking advantage of the copula theory without having to actually estimate the joint distribution is to use copulas as a sophisticated data transformation technique to transform abundances into pseudo-Gaussian data.

### Latent variables

The third popular idea is to model multivariate discrete data using latent variables and push the depdendency back to the latent layer. Latent variables models have recently received increasing attention as they provide a convenient way to model the dependence structure between species. Two specifications of latent variables stand out in community ecology [Warton et al., 2015]: the **Multivariate Generalized Linear Mixed Model (GLMM)** [Ovaskainen et al., 2010, Tingley et al., 2014], and the **Latent Variable Model** (LVM) [Ovaskainen et al., 2016, 2017]. The difference between these models lies in the dimension of their respective random effects: there are as many latent variables as there are species in the GLMM, whereas in the LVM their number is a parameter of the model.

Most methodologies to infer networks from abundance data first use a rationale (data transformation, latent variable modeling, etc.) to solve the problem of network inference in the Gaussian setting. There, they take advantage of the GGM framework to perform network inference using penalized likelihood or tree-based approaches to estimate the precision matrix, from which is finally derived the network.

### Penalized likelihood approaches

There exist two main penalized approaches for the estimation of GGM: the graphical LASSO (glasso) [Friedman et al., 2008], and the neighborhood selection, also called the Meinshausen-Bühlmann approach (MB) [Meinshausen and Bühlmann, 2006]. Both are penalized likelihood approaches which perform a sparse estimation of the precision matrix, either all at once for the glasso or row by row for MB.

### Tree averaging approach

Another GGM inference method considered in this article is the tree averaging approach [Meila and Jordan, 2000], which leverages specific algebraic properties to perform a complete and efficient exploration of the space of spanning tree structures. Note that this approach does not require the GGM Markov faithful property to hold. Each edge is given a posterior probability of being present in the network and those probabilities are thresholded to build the network.

## 7 Supplementary Information

**Table S1:** Species of the cirrhotic guild associated with chronic diseases.

| Species | LIVER<br>Qin et al. [2014] | SCHIZO<br>Zhu et al. [2020] | OB after WL<br>Liu et al. [2017] | AVCD<br>Jie et al. [2017] | CROHN<br>He et al. [2017] |
| --- | --- | --- | --- | --- | --- |
| [Megasphaera] micronuciformis | x | x | x | x |  |
| [Ruminococcus] torques |  |  | x |  | x |
| Bifidobacterium dentium | x | x |  | x |  |
| Campylobacter concisus 2 | x | x |  | x |  |
| Dialister invisus |  | x |  | x |  |
| Haemophilus parainfluenzae | x |  |  |  |  |
| Ligilactobacillus salivarius<br>(ex Lactobacillus) | x |  |  | x |  |
| Limosilactobacillus mucosae<br>(ex Lactobacillus) | x |  |  | x |  |
| Prevotella buccae | x |  |  | x |  |
| Streptococcus anginosus | x | x |  | x |  |
| Streptococcus oralis | x |  |  |  |  |
| Streptococcus parasanguinis | x |  |  | x |  |
| Streptococcus salivarius | x | x |  |  | x |
| Streptococcus thermophilus |  |  |  | x |  |
| Streptococcus vestibularis | x |  |  | x |  |
| Veillonella atypica | x | x |  |  |  |
| Veillonella parvula | x | x |  |  |  |

Among the 23 identified species of the cirrhotic guild, 17 have been found associated together in networks of chronic diseases. The Species column lists these 17 species. Each column named after a chronic disease corresponds to a microbial network shown in the referenced article.

## 8 Supplementary Material

**Supplementary Figure S1:**
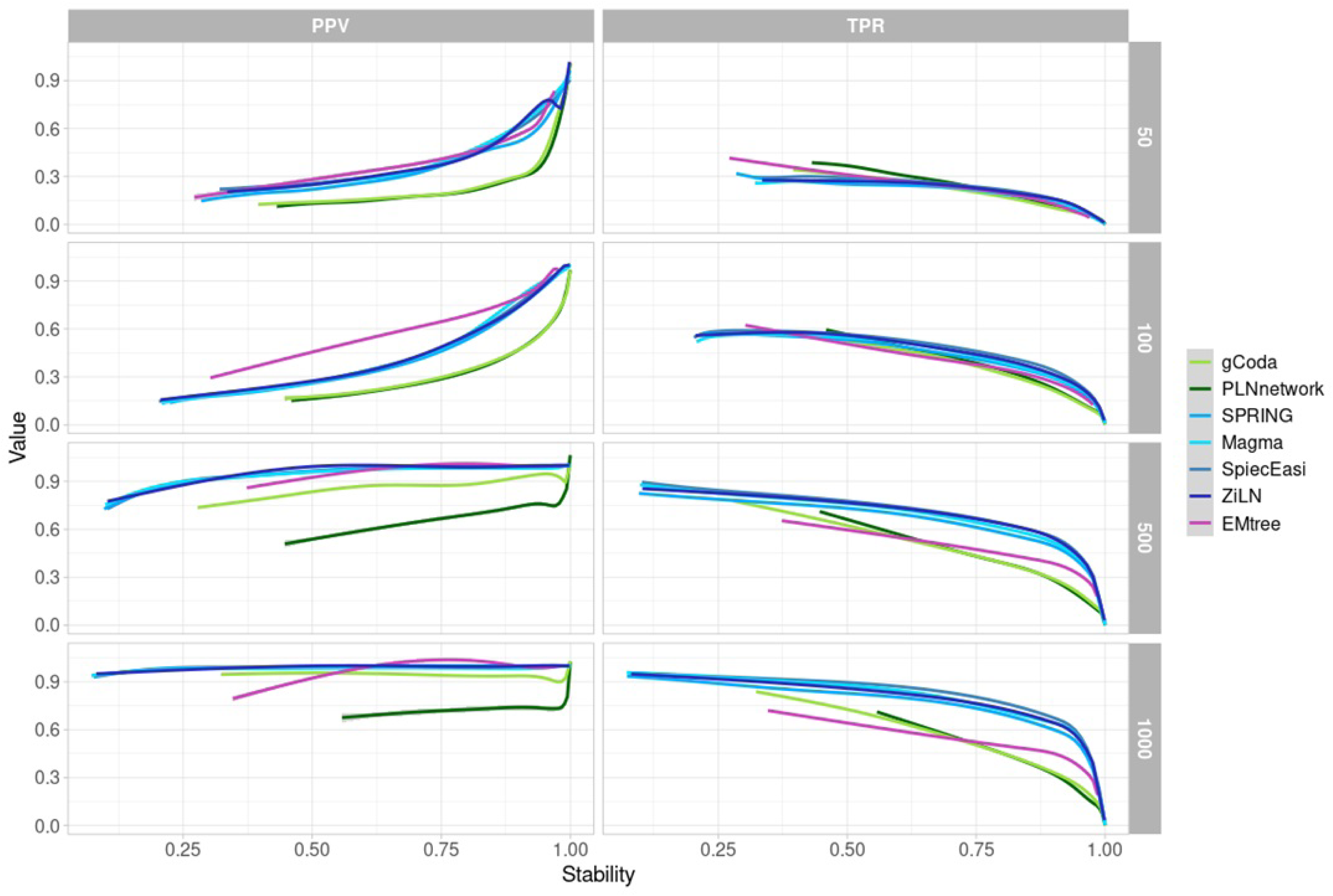
PPV – Stability and TPR – Stability curves of the edge set *E^λ^*(0.90) according to the sample size and inference method. Each point in the curve corresponds to a different value of *λ*.

**Supplementary Figure S2:**
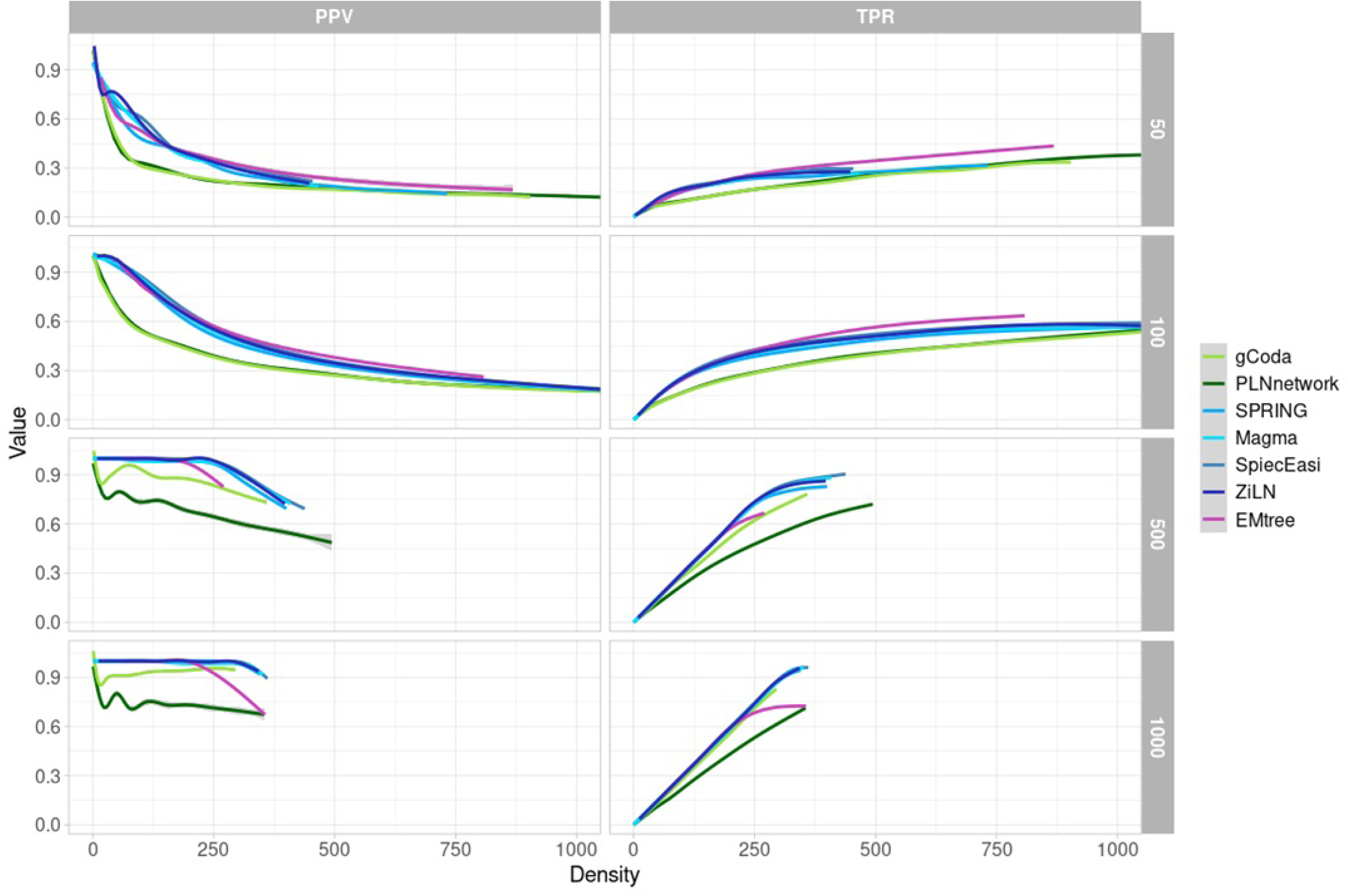
PPV – Density and TPR – Density curves of the edge set *E^λ^*(0.90) according to the sample size and inference method. Each point in the curve corresponds to a different value of *λ*.

**Supplementary Figure S3:**
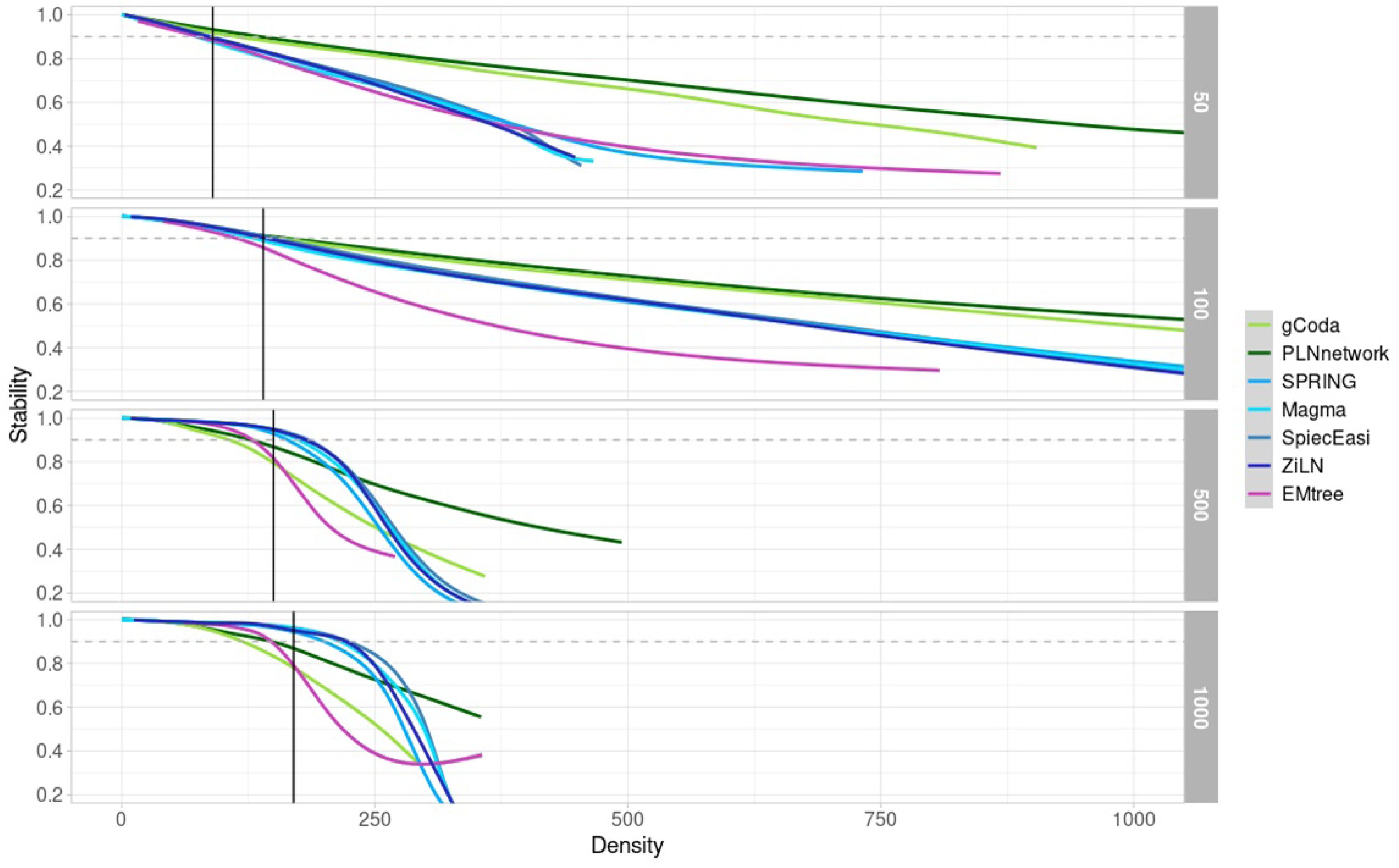
Stability – Density curves of the edge set *E^λ^*(0.90) according to the sample size and inference method. Each point in the curve corresponds to a different value of *λ*. The grey dashed horizontal line represents the target mean stability value (0.90) and the black vertical one, the associated density.

**Supplementary Figure S4:**
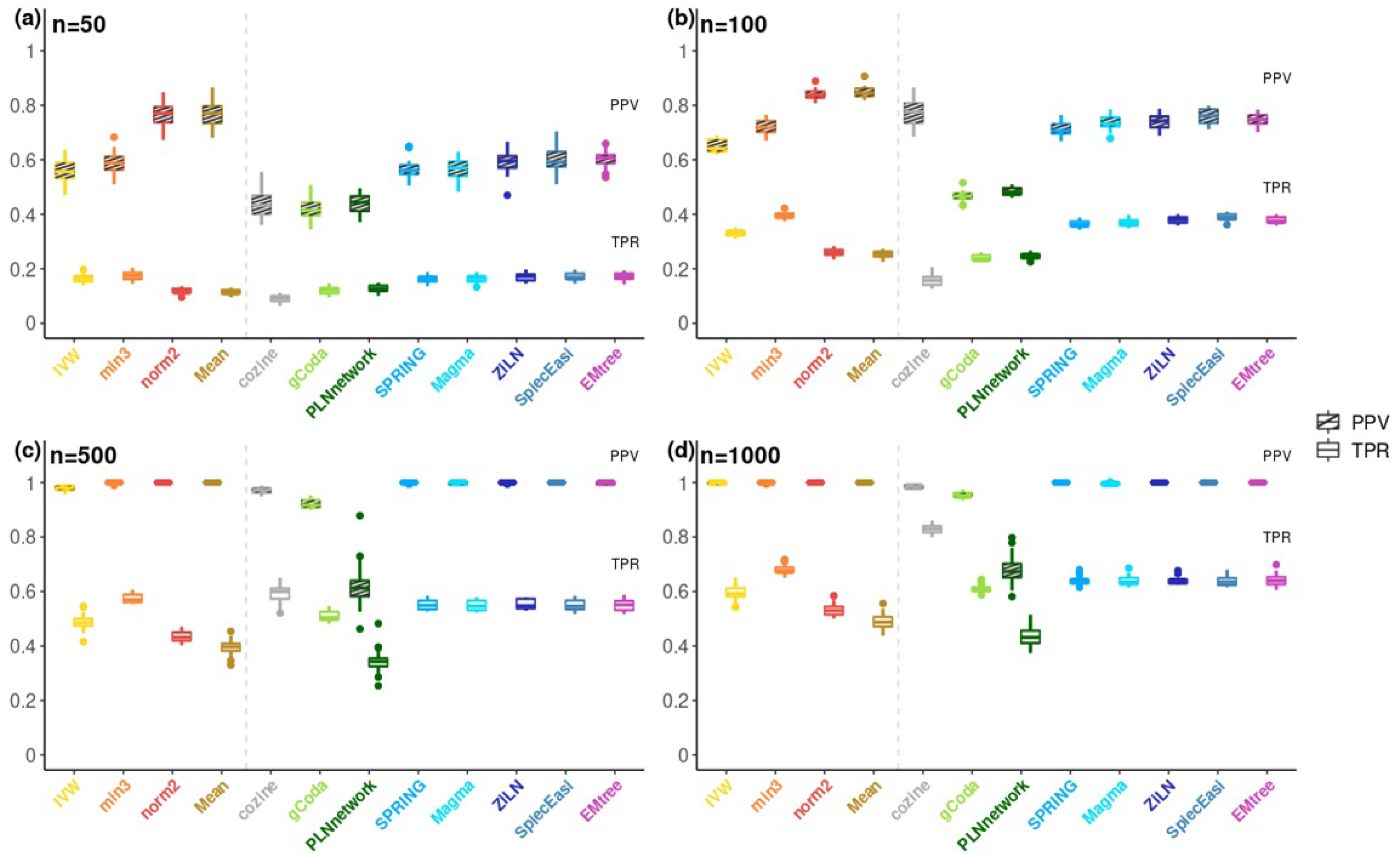
Compared precision (PPV) and recall (TPR) of inference methods and OneNet-* variants when including all 7 methods in the set of methods, for different samples sizes. (a) *n* = 50 (b) *n* = 100 (c) *n* = 500 (d) *n* = 1000.

**Supplementary Figure S5:**
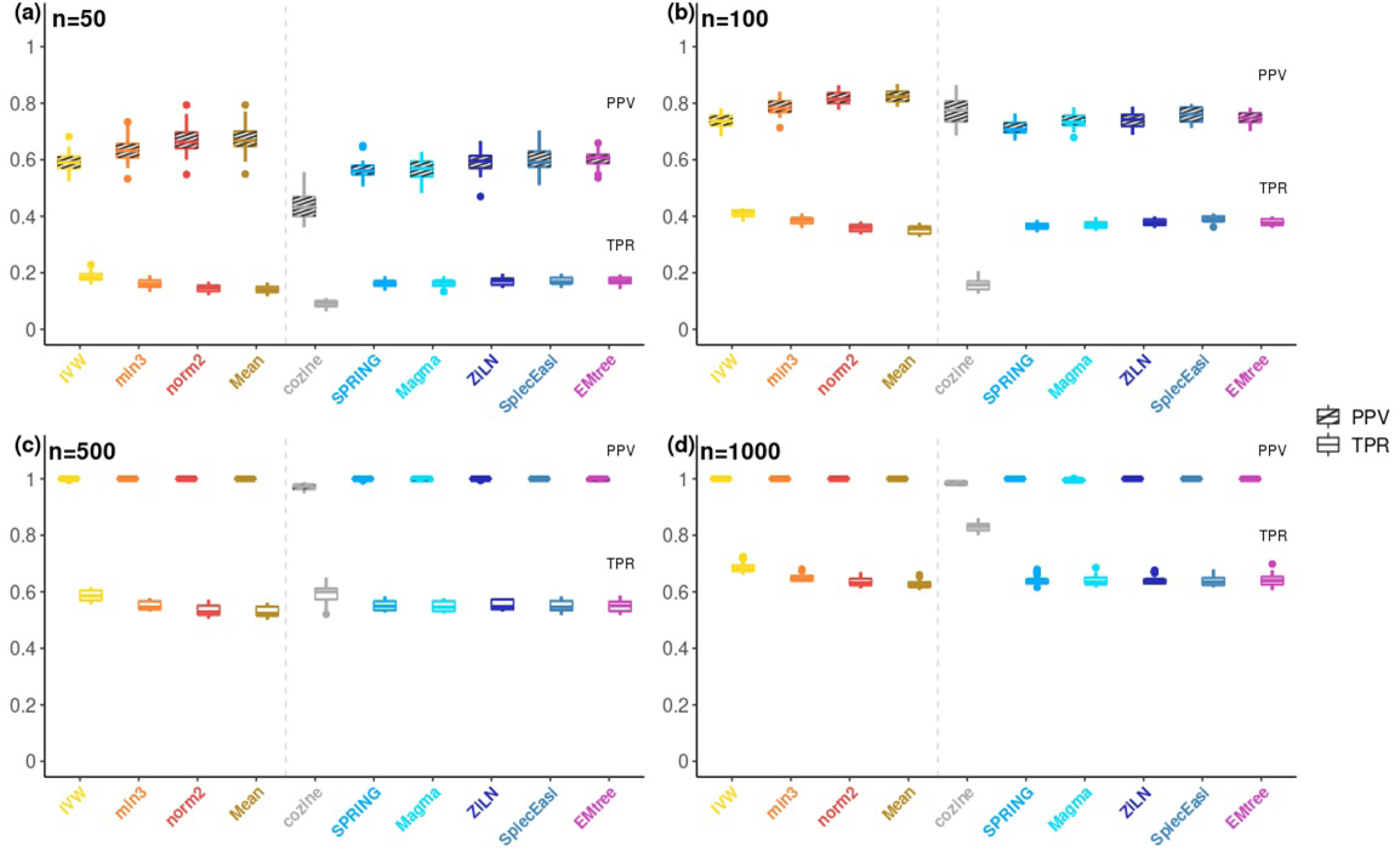
Compared precision (PPV) and recall (TPR) of inference methods and OneNet-* variants after removing glasso-based methods from the set of methods, for different samples sizes. (a) *n* = 50 (b) *n* = 100 (c) *n* = 500 (d) *n* = 1000.

**Supplementary Figure S6:**
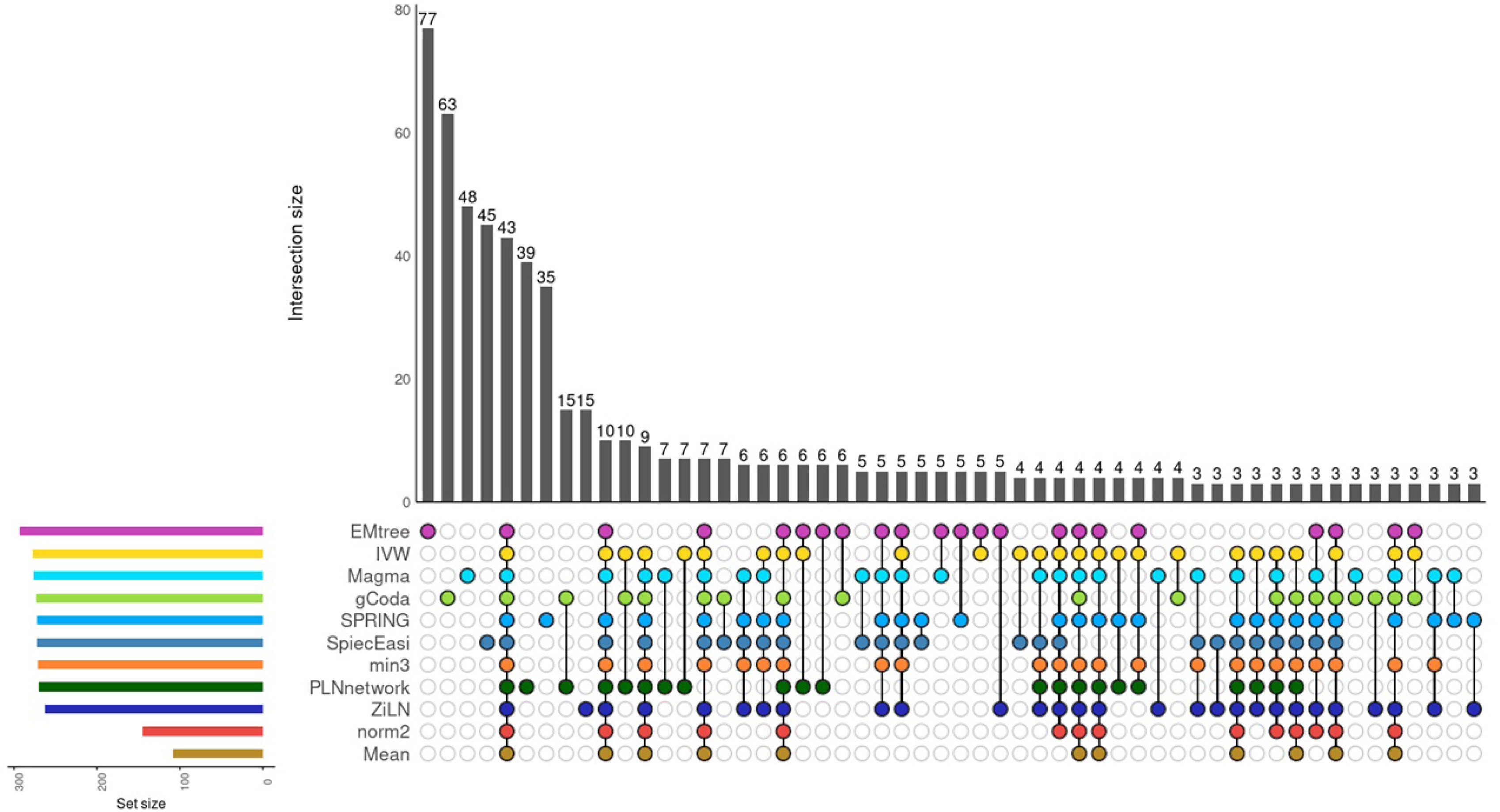
Upset plots of the edges identified by the inference methods and the OneNet-* variants applied to the liver cirrhosis dataset.

**Supplementary Figure S7:**
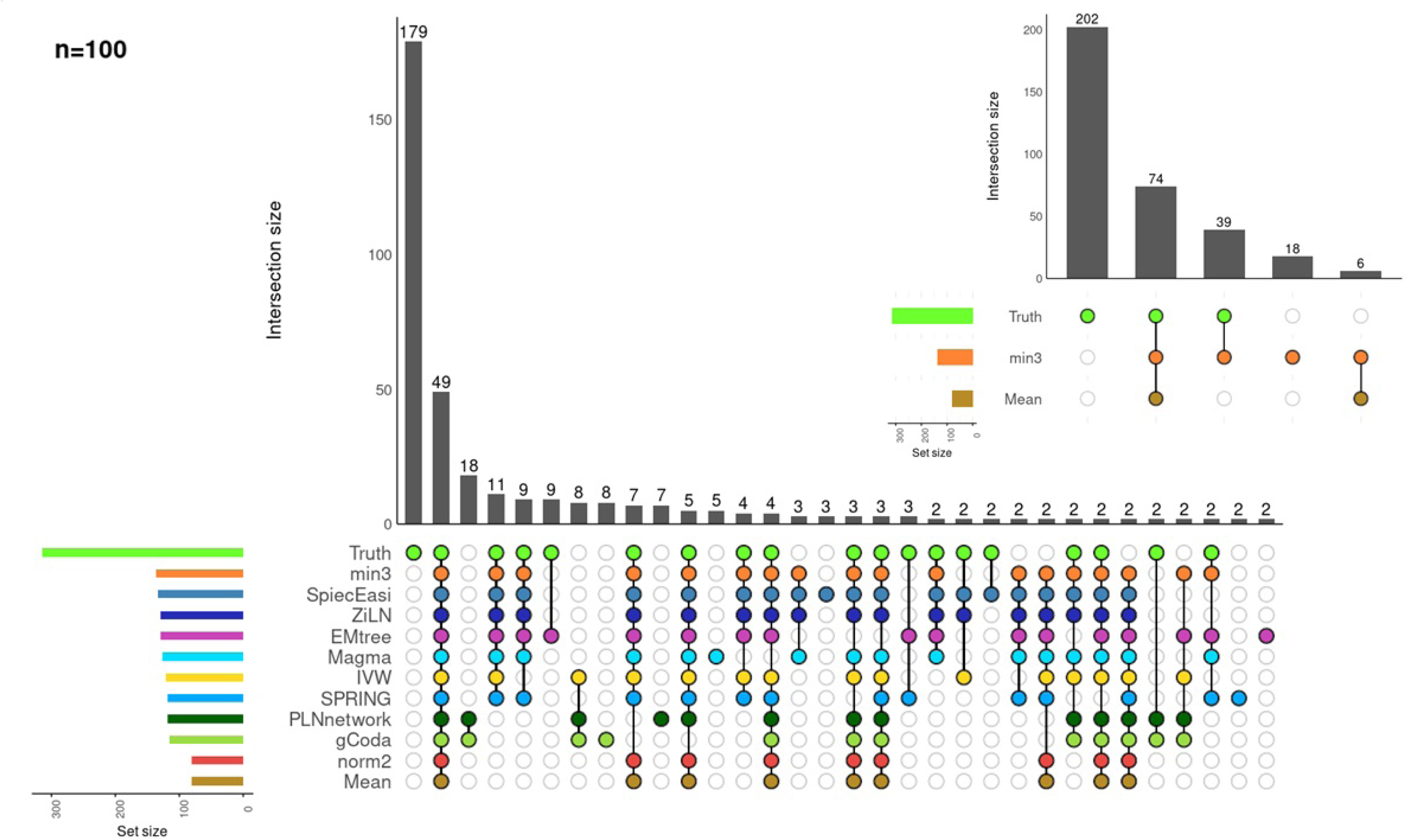
Upset plot of the edges identified by the inference methods, the OneNet-* variants and the ground truth on the synthetic dataset for *n* = 100.

**Supplementary Figure S8:**
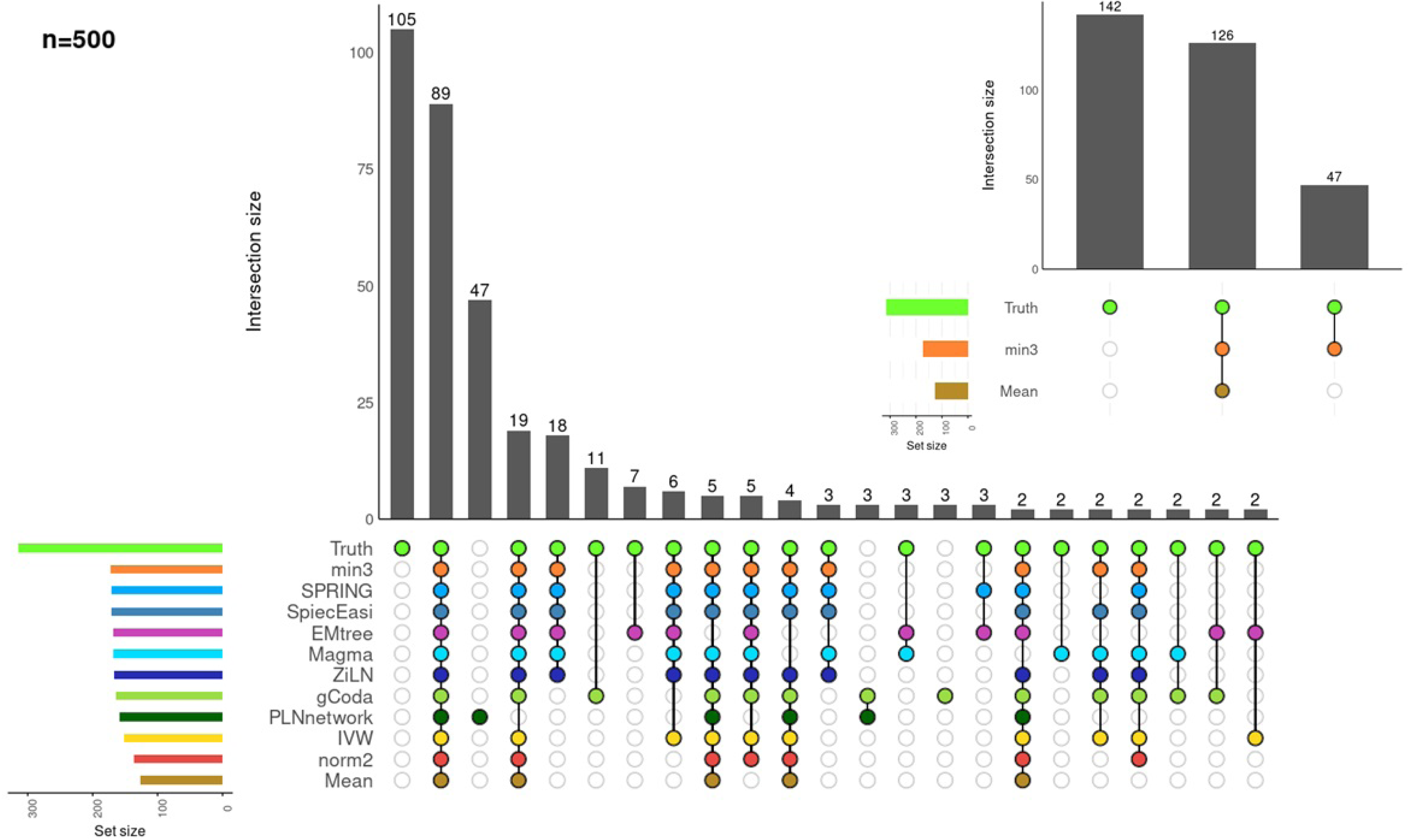
Upset plot of the edges identified by the inference methods, the OneNet-* variants and the ground truth on the synthetic dataset for *n* = 500.

**Supplementary Figure S9:**
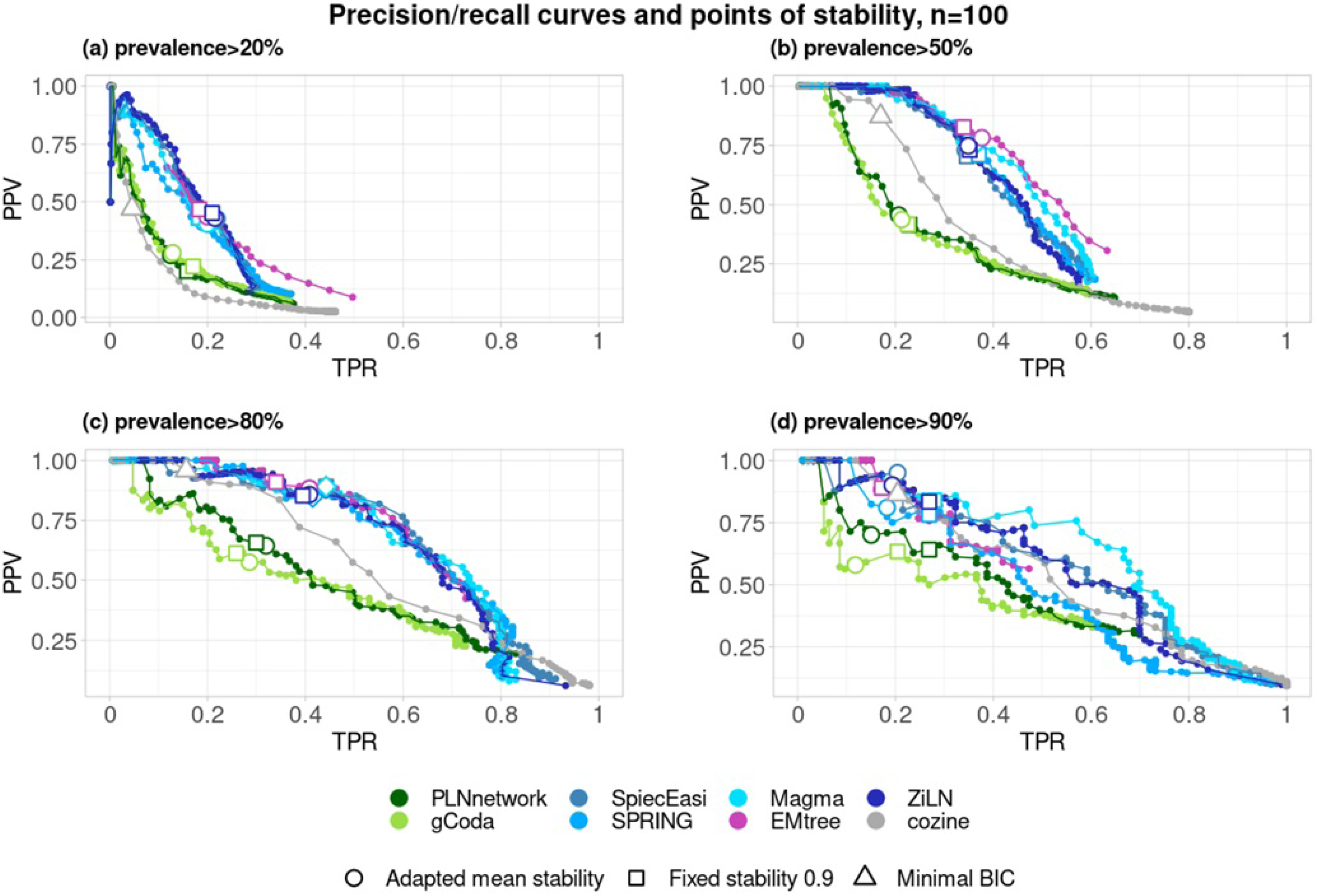
Precision – recall curves of each method and TPR/PPV compromise chosen by stability, mean stability and BIC when *n* = 100 after filtering the dataset to keep only species with prevalence higher than a given threshold. (a)0.20 (b)0.50 (c)0.8 (d)0.9 (see figure 3 for details).

**Supplementary Figure S10:**
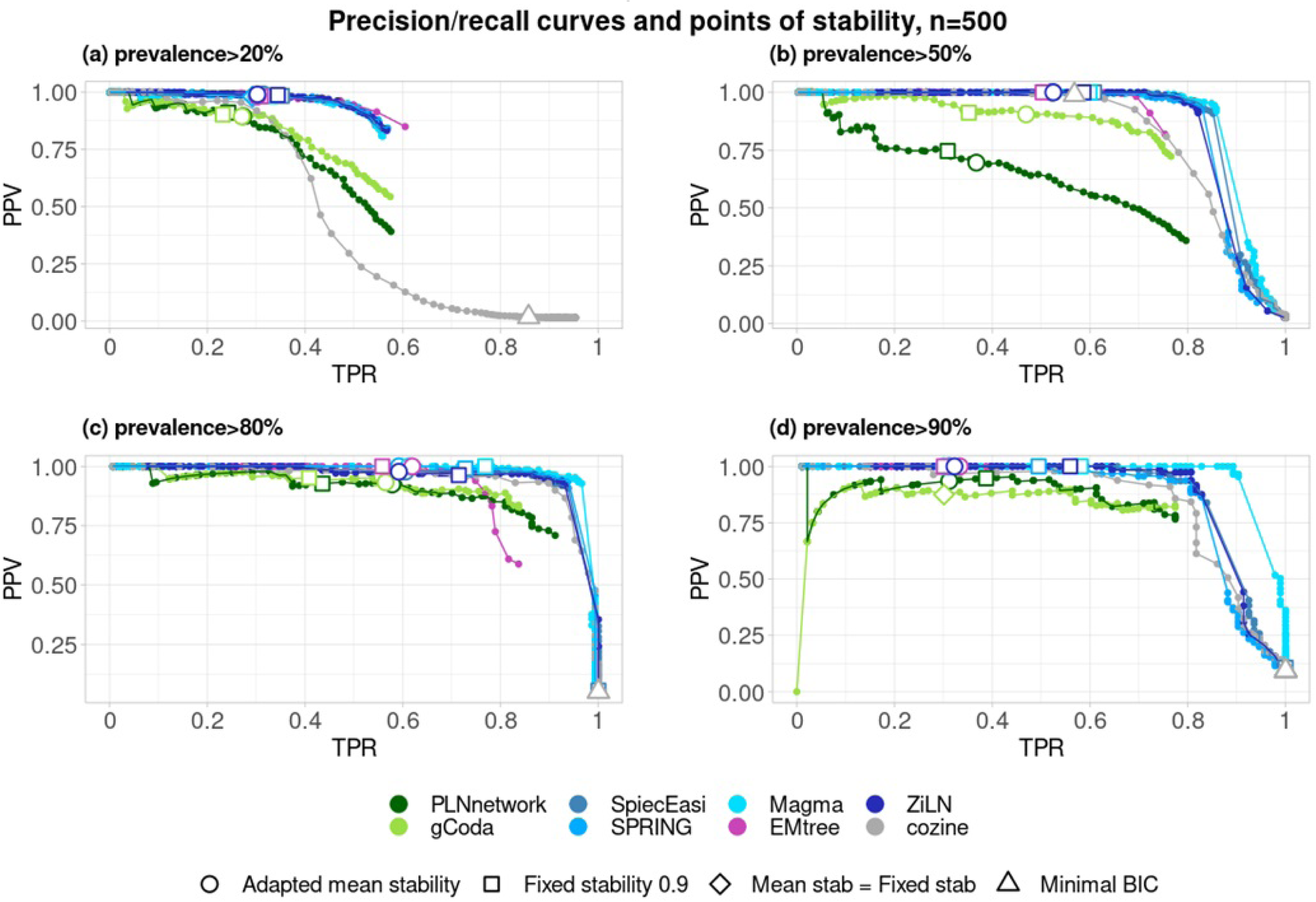
Precision – recall curves of each method and TPR/PPV compromise chosen by stability, mean stability and BIC when *n* = 500 after filtering the dataset to keep only species with prevalence higher than a given threshold. (a)0.20 (b)0.50 (c)0.8 (d)0.9 (see figure 3 for details).

**Supplementary Figure S11:**
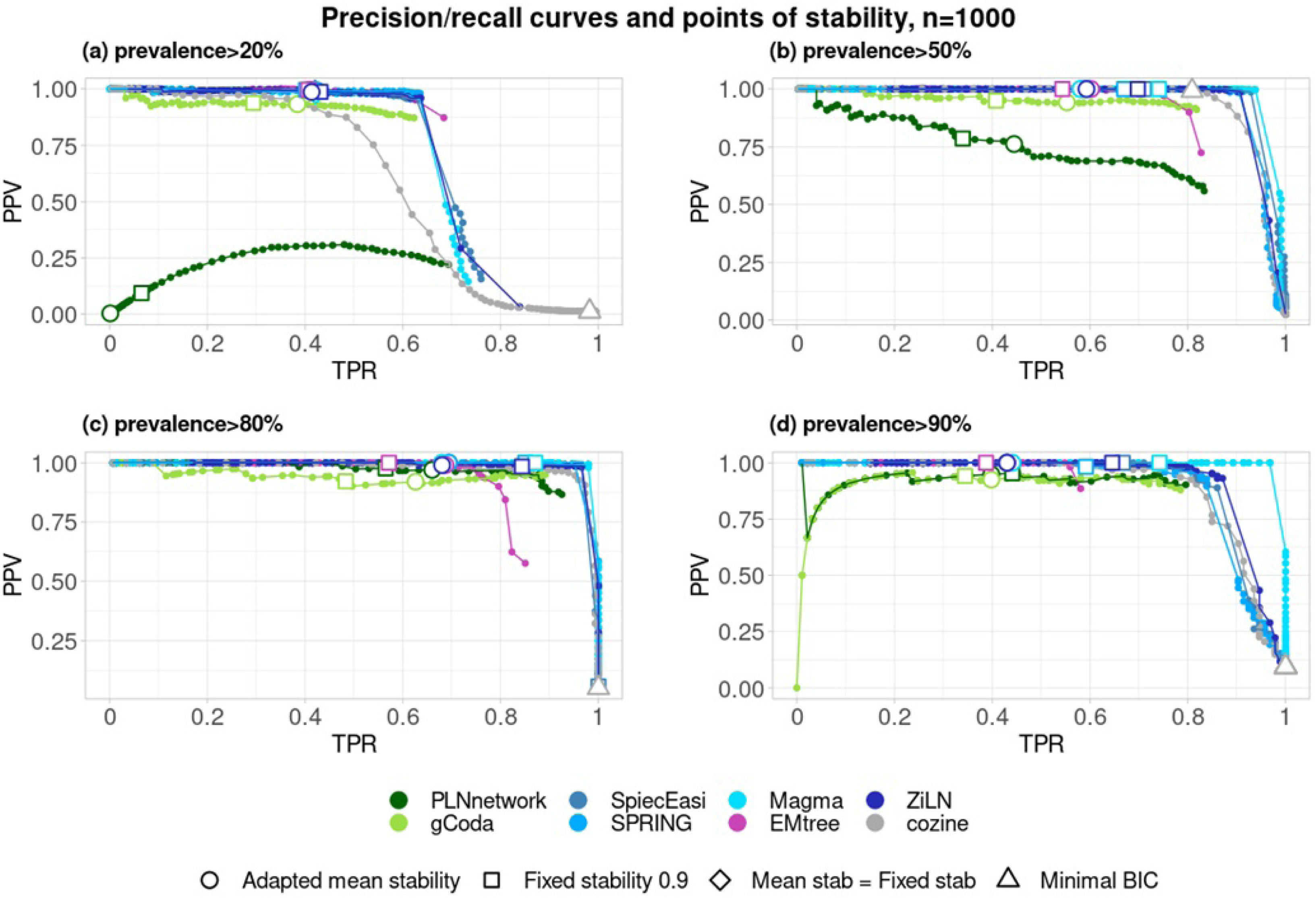
Precision – recall curves of each method and TPR/PPV compromise chosen by stability, mean stability and BIC when *n* = 1000 after filtering the dataset to keep only species with prevalence higher than a given threshold. (a)0.20 (b)0.50 (c)0.8 (d)0.9 (see figure 3 for details).

**Supplementary Figure S12:**
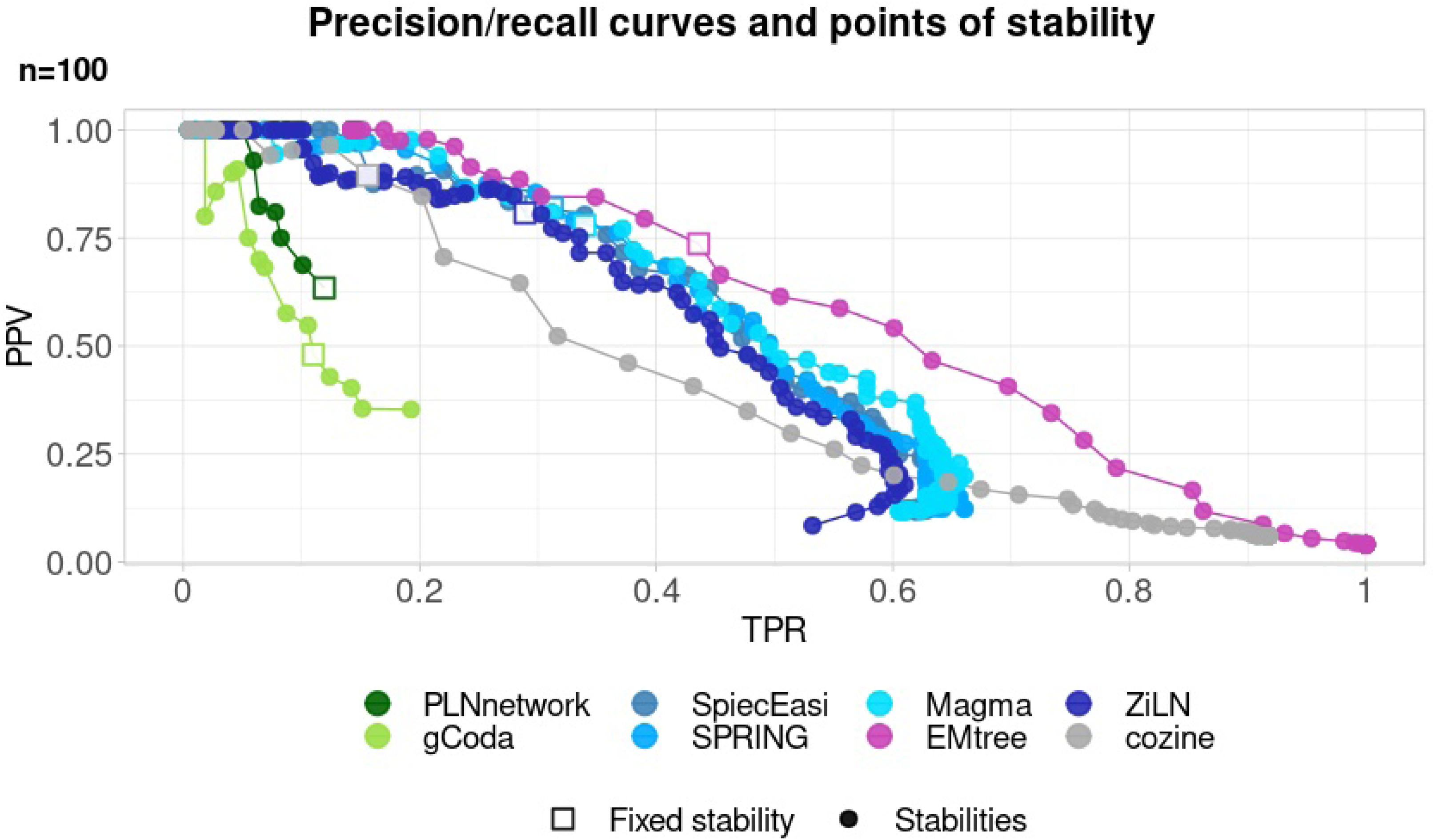
Precision – recall curves of each inference method before equalizing the densities and TPR/PPV value obtained for *λ^∗^* corresponding to default stability criteria shown with a square (*n* = 100). All method have a distinct TPR/PPV compromise and don’t select the same number of edges in the graph.

**Supplementary Figure S13:**
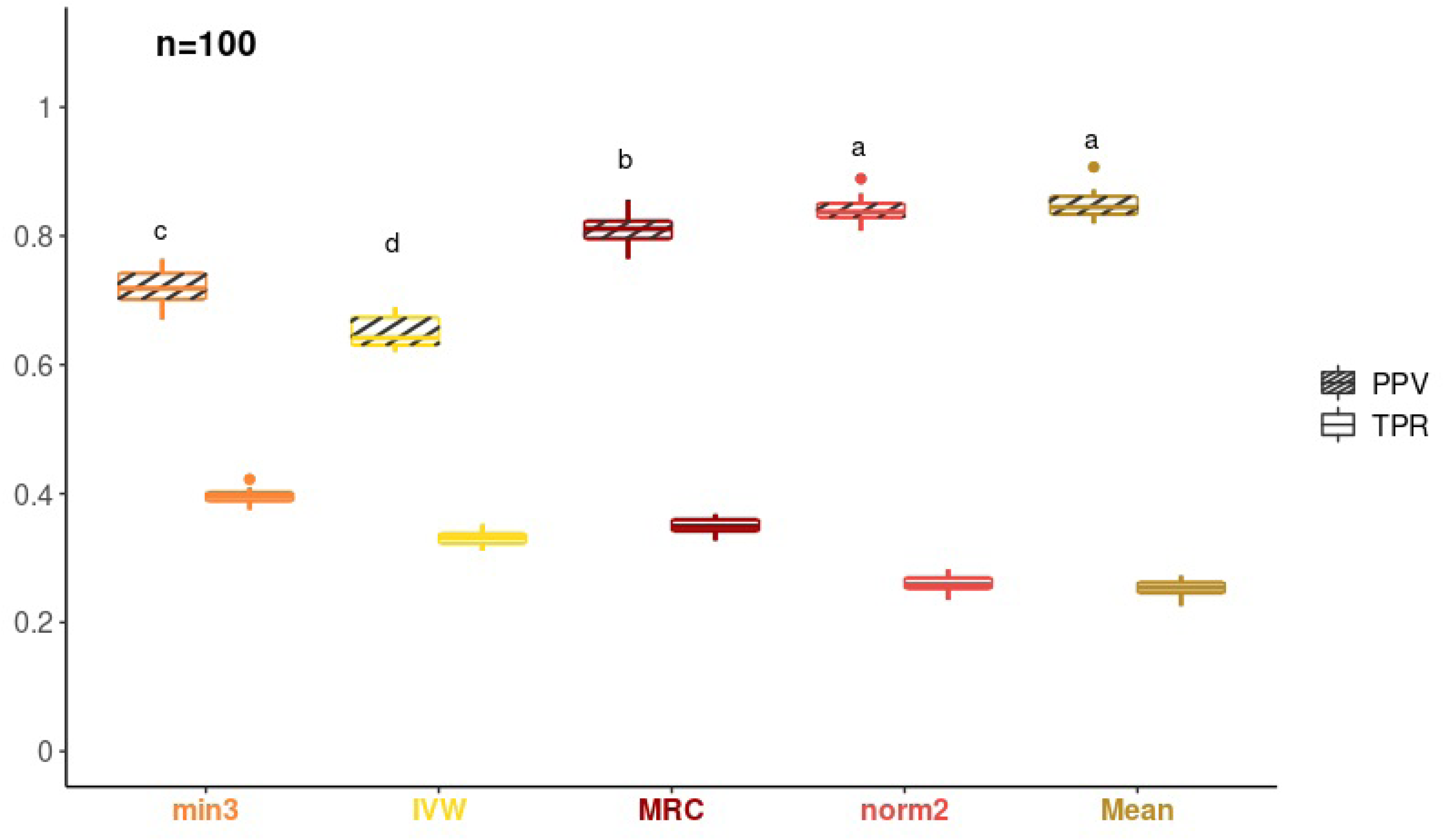
Quality of consensus networks (MRC and OneNet consensus) in terms of PPV/TPR assessed on simulated datasets of size *n* = 100 samples. Striped (resp. no-strip) boxplots show PPV (resp. TPR) values.

**Supplementary Figure S14:**
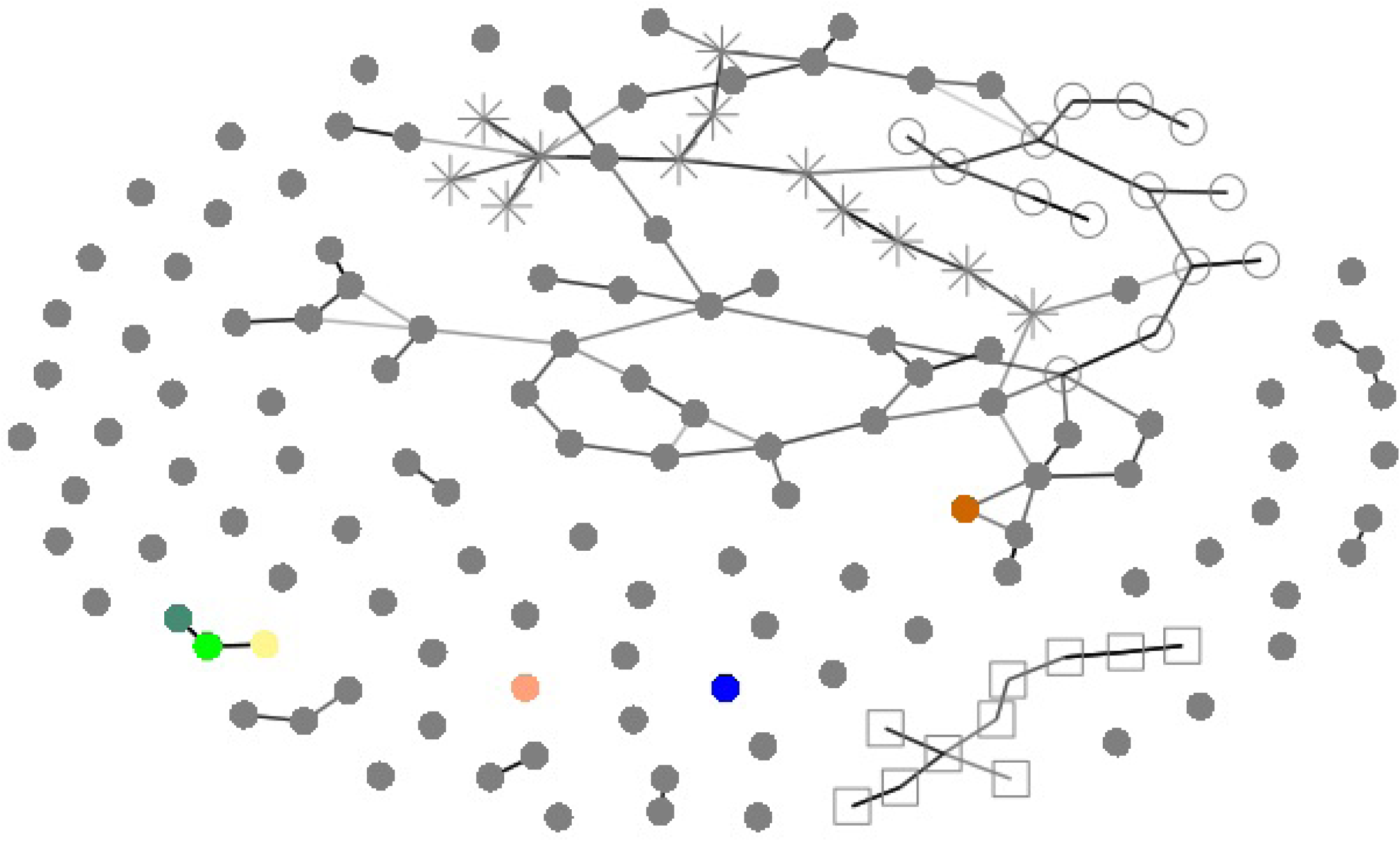
Consensus network inferred on the healthy individuals (n = 102 samples, p = 151 species) from the liver cirrhosis dataset, followed by CORE-clustering algorithm to identify the microbial guilds. The guilds are represented by *,○ and □ and species from the cirrhotic guild are highlighted in color using the same color code as in Fig. 6.

